# Phospho-seq: Integrated, multi-modal profiling of intracellular protein dynamics in single cells

**DOI:** 10.1101/2023.03.27.534442

**Authors:** John D. Blair, Austin Hartman, Fides Zenk, Carol Dalgarno, Barbara Treutlein, Rahul Satija

**Affiliations:** New York Genome Center, New York, NY; New York University, Center for Genomics and Systems Biology, New York, NY; ETH Zurich, Basel, Switzerland

## Abstract

Cell signaling plays a critical role in regulating cellular behavior and fate. While multimodal single-cell sequencing technologies are rapidly advancing, scalable and flexible profiling of cell signaling states alongside other molecular modalities remains challenging. Here we present Phospho-seq, an integrated approach that aims to quantify phosphorylated intracellular and intranuclear proteins, and to connect their activity with cis-regulatory elements and transcriptional targets. We utilize a simplified benchtop antibody conjugation method to create large custom antibody panels for simultaneous protein and scATAC-seq profiling on whole cells, and integrate this information with scRNA-seq datasets via bridge integration. We apply our workflow to cell lines, induced pluripotent stem cells, and 3-month-old brain organoids to demonstrate its broad applicability. We demonstrate that Phospho-seq can define cellular states and trajectories, reconstruct gene regulatory relationships, and characterize the causes and consequences of heterogeneous cell signaling in neurodevelopment.

## Introduction

The ability to carefully regulate responses to external and internal signals is essential for proper cellular function. Maintaining precise control of signaling networks enables cells to achieve homeostasis in response to environmental changes and to progress through key developmental and functional transitions ^1^. Signal transduction links the activation of signaling pathways to downstream changes in cellular chromatin ^2^, transcription ^3^, and translation ^4^, and is primarily regulated by changes in post-translational modifications (PTMs) ^5^. For example, changes in phosphorylation can significantly alter receptor kinetics ^6^, enzymatic function ^7^, and transcription factor localization and activity ^8^ via conformational changes ^9^.

Cell signaling pathway activity is known to vary throughout neurodevelopment, and when disrupted, it can have significant effects on cell fate decisions. For instance, uncontrolled mTOR pathway activity caused by the loss of the upstream regulator TSC2 leads to a significant increase in astrogliosis, which is a hallmark of the neurodevelopmental disorder, tuberous sclerosis ^10^. Additionally, changes in Wnt signaling have been associated with autism spectrum disorder ^11^. In order to gain a deeper understanding of the causes and consequences of aberrant signaling, a crucial challenge is to connect heterogeneity in the activation of signaling pathways with broader changes in molecular state. Therefore, methods that can accurately measure and quantify phosphorylation states at single-cell resolution alongside additional molecular modalities offer substantial promise to improve our understanding of cellular activity and function.

Phospho-proteins are typically quantified in single-cells using antibody-based methods including immunocytochemistry, flow cytometry ^12^, or CyTOF ^13^, which can be multiplexed to detect dozens of targets. Single-cell sequencing technologies offer an exciting opportunity to build upon these approaches, particularly with the recent introduction of “multiomic” technologies that enable the quantification of multiple modalities of information within the same cell ^14–21^. For example, CITE-seq ^15^, REAP-seq ^19^, DOGMA-seq ^18^, and TEA-seq ^20^ all stain cells with large panels of oligonucleotide-tagged antibodies against cell surface proteins in order to quantify cellular immunophenotypes alongside cellular transcriptomes, chromatin accessibility profiles, or both. While powerful, these technologies focus exclusively on the profiling of cell surface proteins. They are therefore widely applied to analyze hematopoietic samples, where well-characterized panels of cell surface proteins are associated with distinct cell states, but have limited utility in other contexts, including neurodevelopment.

Multiple pioneering approaches have built upon these methods, aiming to utilize single-cell multimodal technologies to profile intracellular and intranuclear proteins in diverse biological contexts. These include ASAP-seq ^18^, which introduces a set of fixation and permeabilization conditions that are compatible with chromatin accessibility profiling in whole cells. Additionally, inCITE-seq ^21^ and NEAT-seq ^16^ utilize specialized approaches for intranuclear protein profiling, which significantly reduce background signal originating from non-specific electrostatic interactions between oligonucleotide-conjugated antibodies and charged cellular components. While each of these methods addresses key challenges, they perform profiling of small (3-7) intracellular panels due to a reliance on commercially conjugated antibodies. QuRIE-seq aims to profile larger panels, but requires custom instrumentation and is not compatible with primary cell samples ^17^. Lastly, while NEAT-seq utilizes the 10x Multiome kit for trimodal nuclear profiling, no existing approaches can quantify intracellular proteins (including phosphorylation states), transcriptional output, and chromatin profiling in the same biological system.

Here we present Phospho-seq, a multi-modal single-cell workflow for quantifying cell signaling via phosphorylated cytoplasmic and nuclear proteins in conjunction with chromatin accessibility and, through bridge integration ^22^, gene expression levels. By optimizing a broadly accessible antibody conjugation strategy ^23^, we designed a custom panel to profile 64 intracellular proteins (including 20 phospho-states), and applied this workflow to 3-month old brain organoids. We used the trimodal chromatin, RNA and protein data to discover novel transcription factor-cis-regulatory element-gene associations including telencephalic and diencephalic lineage specific relationships. We further used the phosphorylated protein data to identify lineage-specific patterns of WNT and MAPK/ERK signaling, and to link these differences to upstream and downstream molecular networks. Overall, we demonstrate the utility of Phospho-seq for the high throughput discovery of interactions between cell signaling, gene regulation and gene expression in neural tissue, opening the door for future discoveries in other cell types and tissues.

## Results

### Benchtop conjugation enables antibody panel customization in Phospho-seq

In developing Phospho-seq, we aimed to create a user-friendly, single-cell method to quantify proteins from the cell surface, cytoplasm and the nucleus alongside additional molecular modalities. To maximize Phospho-seq’s utility, we aimed for the assay to 1) allow for maximum customizability by the user for which proteins to quantify, 2) rely only upon commercially available reagents and equipment, and 3) maintain the sensitivity and specificity of single-modality assays. Towards these goals, Phospho-seq combines aspects of previously established single-cell protein and chromatin accessibility quantification methods ^16,18^. In brief, cells are dissociated into single-cells, fixed, permeabilized, hashed, stained for intracellular proteins with self-conjugated DNA-bound antibodies, and run through the 10X Genomics single-cell ATAC-seq protocol (Fig 1A and Methods).

**Figure 1:**
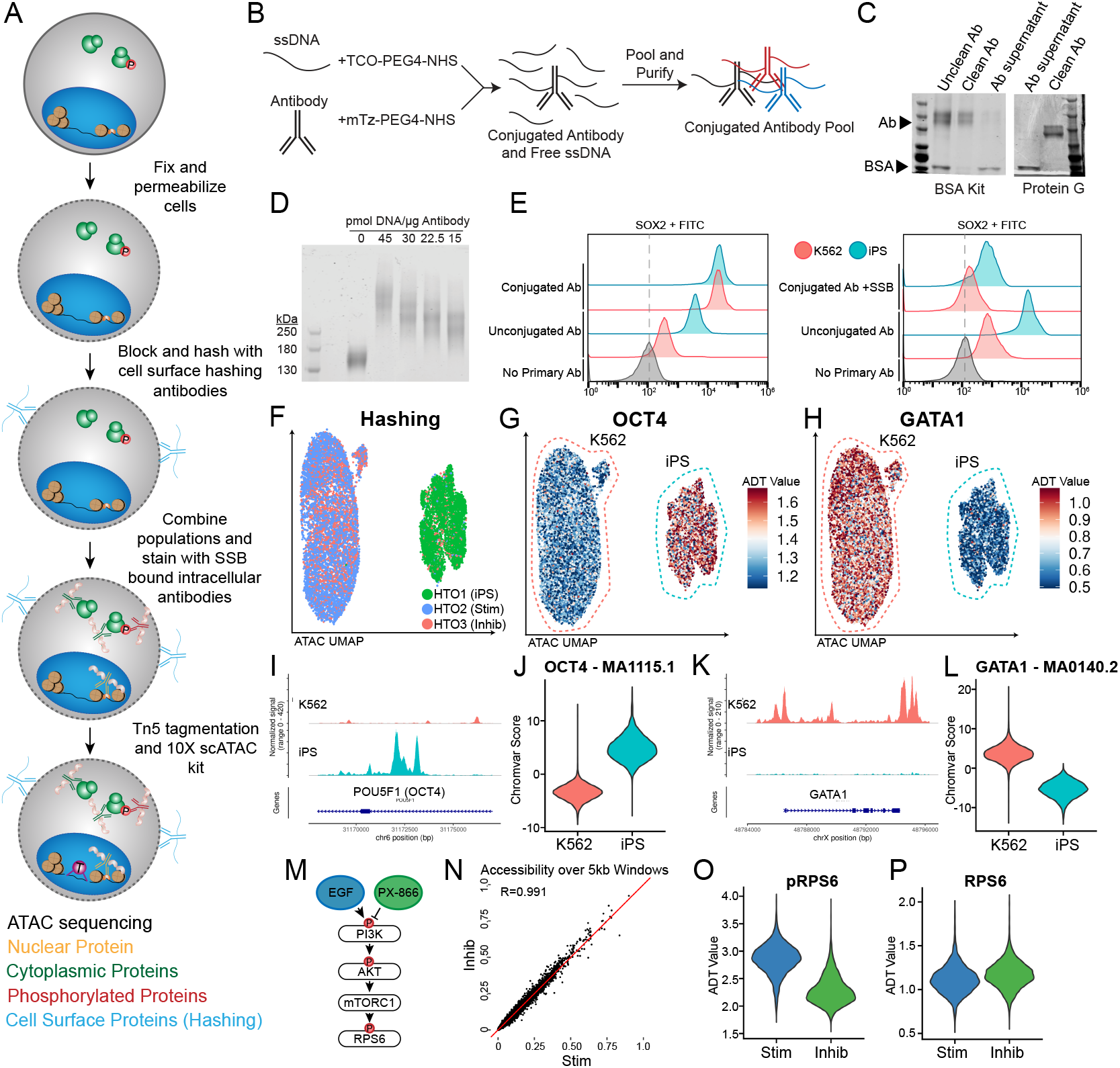
Phospho-seq procedure and pilot experiment. **a)** Schematic of Phospho-seq workflow. **b)** Schematic of antibody conjugation procedure. **c)** Protein gel results of two antibody purification methods using the Abcam BSA Removal Kit (left panel) and Promega Magne Protein G beads (right panel). **d)** Protein gel of mTz-PEG4-NHS labeled antibodies incubated with different quantities of TCO-PEG4-NHS labeled ssDNA tags. **e)** Flow cytometry plots of K562 and iPS cells stained with unconjugated and conjugated SOX2 antibodies (left panel) and unconjugated and conjugated + single-stranded DNA binding protein SOX2 antibodies (right panel) with unstained controls. **f)** UMAP representation from scATAC-Seq of K562 and iPS cells colored by demultiplexed HTOs assigned to each cell. **g)** UMAP representation of K562 and iPS cells colored by normalized ADT values for OCT4. **h)** UMAP representation of K562 and iPS cells colored by normalized ADT values for GATA1. **i)** Coverage plot of chromatin accessibility of K562 and iPS cells at the POU5F1 (OCT4) genomic locus. **j)** Violin plot of chromVAR scores for the OCT4 TF binding motif (MA1115.1) in K562 and iPS cells. **k)** Coverage plot of chromatin accessibility of K562 and iPS cells at the GATA1 genomic locus. **l)** Violin plot of chromVAR scores for the GATA1 TF binding motif (MA0140.2) in K562 and iPS cells **m)** Schematic of PI3K/AKT/mTORC1 pathway activation and repression paradigm used in this experiment. **n)** Scatter plot of pseudobulked chromatin accessibility data in 5 kb windows across the genome comparing inhibited K562 cells with stimulated K562 cells. Red line indicates perfect correlation between the two conditions. **o)** Violin plot of normalized pRPS6 values in stimulated (Stim) and inhibited (Inhib) K562 cells. **p)** Violin plot of normalized RPS6 values in stimulated (Stim) and inhibited (Inhib) K562 cells.

A major limitation of large-scale sequencing-based intracellular profiling is the lack of commercially available oligonucleotide-tagged antibodies. Premade antibody panels are specific for hematopoietic surface antigens ^24^, while custom commercial antibody conjugation is often prohibitively expensive, especially for larger panels. To overcome this issue, we optimized a simple benchtop click-chemistry-based conjugation protocol ^23^ (Fig 1B) to generate panels of uniquely-indexed DNA-bound antibodies. This approach is scalable, cost-effective (∼$8/conjugation), and compatible with unconjugated commercial antibodies that are routinely used for immunofluorescence or flow cytometry, even those with carrier proteins (Fig 1C). Phospho-seq is compatible with antibodies conjugated to different oligonucleotide sequences, including the widely used TotalSeq-A and TotalSeq-B sequences, through the use of a “bridge oligonucleotide” for capture on the gel beads ^18^.

During the optimization of our conjugation protocol, we found that the ratio of antibody to oligonucleotide and post-conjugation purification steps were crucial for minimizing nonspecific binding while maintaining a high recovery yield. Through titration experiments, we determined that adding 15 pmol of oligonucleotide per μg of antibody (equivalent to 2-4 copies of oligonucleotide per antibody molecule) was optimal (Fig 1D). For post-conjugation purification, we found that two steps: an initial precipitation step using 40% ammonium sulfate ^25^, followed by 5-7 washes through a 50 kDa molecular weight cut-off (MWCO) filter, were necessary to reduce unbound oligonucleotide while preserving antibody yield (Fig S1A).

We utilized flow cytometry experiments on whole cells to optimize and confirm that our Phospho-seq procedure was capable of retaining cell-to-cell differences in intracellular antibody levels. We first confirmed that light fixation (1% formaldehyde) and gentle detergent-based permeabilization (0.1% NP-40), as suggested by ASAP-seq ^18^, achieved the necessary balance between maintaining the structural integrity of the cell membrane while allowing the flow of antibodies into the nucleus and cytoplasm. We also found that the addition of single-stranded DNA binding protein (SSB) to our antibody pool before staining, as pioneered for nuclear profiling with NEAT-seq ^16^, was essential to reduce background signal for oligonucleotide-conjugated antibodies. We therefore combined these approaches, and utilized flow cytometry for an oligonucleotide-conjugated antibody against SOX2 to successfully quantify clear expression differences in heterogeneous mixtures of whole cells (Fig 1E).

### Phospho-seq simultaneously quantifies phosphorylated, cytoplasmic and nuclear proteins

To evaluate the full Phospho-seq workflow on whole cells, we first tested a small panel of both nuclear and cytoplasmic proteins on a heterogeneous cell line mixture. Our panel included antibodies against the transcription factors OCT4, SOX2, and GATA1, which are expected to be differentially expressed between K562 cells and induced pluripotent stem cells (iPSCs). We also aimed to quantify phosphorylated ribosomal protein S6 (pRPS6) expression, a readout of PI3K/AKT/mTOR pathway activation, along with antibodies quantifying total RPS6 levels. We exposed cells to either epidermal growth factor for 1 hour (EGF, activating) or PX-866 for 4 hours (inhibiting) to modulate pathway activation, and utilized cell surface ‘hashing’ antibodies to perform multiplexed profiling of cells in resting, activated, or inhibited conditions.

We found that Phospho-seq was able to accurately quantify intranuclear, intracellular (including phosphorylated), and cell surface proteins alongside chromatin accessibility. Dimensional reduction of scATAC-seq profiles successfully discriminated the two cell lines and was concordant with the cell hashing-based demultiplexing. (Fig 1F and S1B,C). Antibody-derived tags (ADTs) for the defining canonical transcription factors OCT4, SOX2 (iPSCs) and GATA1 (K562) showed strong, significant differences between the two cell types (Fig 1G,H and Fig S1D). These profiles were also concordant with the chromatin accessibility landscape at each protein’s genomic locus (Fig 1I,K and S1E) and genome-wide transcription factor motif activity estimates as quantified by chromVAR ^26^ (Fig 1J,L and S1F).

Moreover, while changes in PI3K/AKT/mTOR pathway (Fig 1M) activity did not drive genome-wide changes in chromatin accessibility (Fig 1N), we observed a clear increase in pRPS6 levels within stimulated cells compared to inhibited cells (Fig 1O). These results were not echoed in total RPS6 levels (Fig 1P), and reflect a biological context where phosphorylation measurements are highly informative for distinguishing cellular states even when genome-wide modalities cannot. We also observed higher phosphorylation of the nuclear transcription factor STAT3 in iPSCs compared to K562 cells, and found that only pSTAT3 levels (as opposed to total protein levels) correlated with STAT3 transcription factor activities (Fig S1G). We conclude that we can therefore measure phosphorylation states for both intracellular and intranuclear activities, and that phosphorylated protein levels can more accurately reflect cellular state and transcription factor activities compared to total protein levels.

We further observed that Phospho-seq was capable of quantifying more subtle differences, even within the same cell type. Epigenetic differences between iPS donor cell lines are frequently observed and often lead to biased differentiation tied to cell signaling differences ^27^. In our dataset, we observe the segregation of the three iPSC donors used in this experiment based on chromatin accessibility (Fig S1H) as well as an enrichment of pRPS6 signal in one donor in particular (Fig S1I). These types of observations may be highly informative when assessing the differentiation capacity of different iPSC lines.

### Intracellular protein staining highlights cell type differences in human brain organoids

We next applied Phospho-seq to human iPSC-derived brain organoid models, which have provided valuable insights into early neurodevelopmental processes that are otherwise difficult to study in humans ^28,29^. We first designed and conjugated a custom panel of 64 antibodies comprising both of neurodevelopmentally relevant transcription factors, as well as an expanded set of 40 paired cell-signaling antibodies (Fig 2A and Table S1). While previous studies have profiled chromatin and transcriptomic modalities from these models ^29,30^, we reasoned that our Phospho-seq panel could illuminate relationships between cell signaling pathways, transcription factors, and gene regulatory elements. We separately generated 3-month-old organoids from four iPS donors, using a well-established protocol for unguided brain organoid differentiation (Fig 2B) ^29,31^. We performed cell hashing for doublet detection ^32^ and used genetic profiling ^33^ for donor demultiplexing, subsequently profiling 9,028 cells using Phospho-seq (Fig S2A).

**Figure 2:**
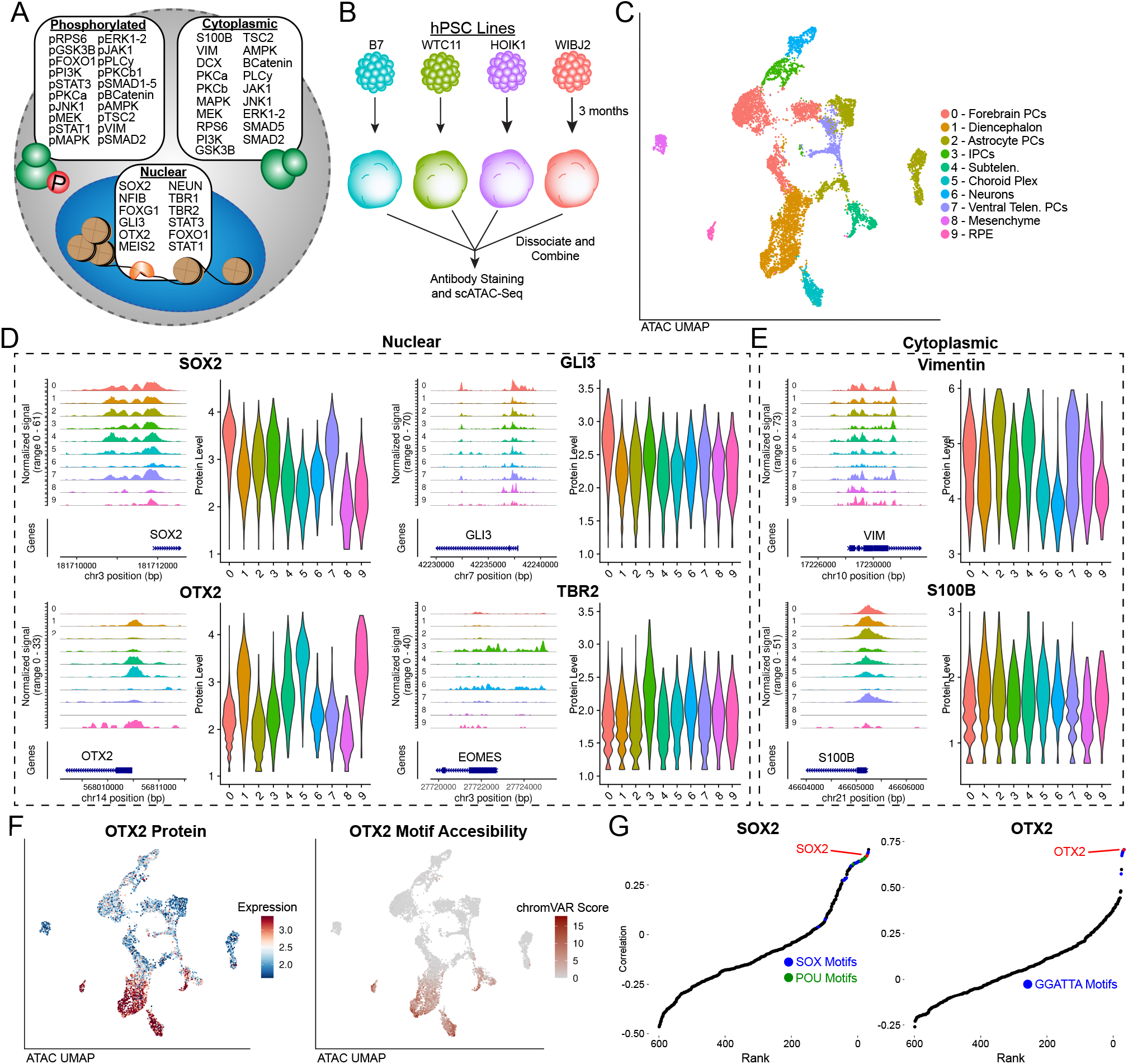
Phospho-seq on brain organoids. **a)** Schematic of antibody panel used in brain organoid Phospho-seq experiment and the cellular compartment of the target protein. **b)** Schematic of brain organoid differentiation. **c)** UMAP representation of cells and cell type assignments based on the ATAC-seq modality in Phospho-seq. **d)** Coverage and violin plots of the gene promoters and protein levels respectively of four nuclear proteins included in the Phospho-seq panel (SOX2,GLI3,OTX2 and TBR2), color and order are same as in **(c). e)** Coverage and violin plots of the gene promoters and protein levels respectively of two cytoplasmic proteins included in the Phospho-seq panel (Vimentin and S100B), color and order are same as in **(c). f)** UMAP representation of cells colored by normalized ADT values for OTX2 (left panel), UMAP representation of cells colored by chromVAR scores for OTX2 motif accessibility. **g)** Rank-correlation plots for transcription factor motifs vs. SOX2 (left panel) and OTX2 (right panel) with the TF motif of interest indicated in red.

Utilizing dimensionality reduction and unsupervised clustering, we identified 15 clusters representing heterogeneity both in neurodevelopmental lineage and donor identity (Fig S2B). While we could assign broad cell type labels using gene activity scores calculated from chromatin accessibility within gene bodies and promoters ^34^ (Fig 2C and S2C), more fine-grained annotation is challenging for chromatin accessibility measurements in the absence of transcriptomic data. We found that ADTs were heterogeneously detected across differently assigned cell types, identifying markers of forebrain progenitors (SOX2,GLI3), diencephalic cells (OTX2), progenitors and glia (VIM,S100B), and intermediate progenitors (TBR2). We observed that 45 out of 64 proteins exhibited differential expression between at least one pair of clusters (Fig S2D), and found that elevated protein marker expression was further supported by enriched chromatin accessibility patterns in the same cell types for both intranuclear and intracellular proteins (Fig 2D,E). We observed clear concordance between single-cell transcription factor ‘activities’ as measured by chromVAR, and transcription factor levels measured by Phospho-seq (Fig 2F and S2E). Strikingly, when correlating the measured protein levels of OTX2 with all 633 chromVAR motifs, the OTX2 motif itself was the second highest hit (Fig 2G), while the other top hits had the same core motif. We observed similar results for SOX2 (Fig 2G), and concluded that Phospho-seq data can help to identify and discover *bona fide* links between cellular protein levels and DNA sequences that influence gene expression.

### High Quality RNA expression data is incorporated into Phospho-seq through bridge integration

Our Phospho-seq protocol utilizes the 10x scATAC-seq kit to generate scalable and high-quality chromatin accessibility profiles, but lacks transcriptomic measurements which are highly valuable for fine-grained cell annotation and gene regulatory network reconstruction. In considering how to add these measurements to our analysis, we generated two additional datasets on the same single-cell suspension of 3-month old brain organoids: scRNA-seq (3’ 10x Chromium), and scRNA-seq+scATAC-seq (10x Multiome). We followed standard manufacturer’s protocols for both experiments, and observed that obtaining simultaneous measurements from the multiomic dataset was associated with a significant reduction in data quality compared to single-modality assays. Even after controlling for sequencing saturation, multiomic profiles exhibited a more than five-fold reduction in sensitivity for detected transcripts (Table S3). We conclude that further modifying the 10x Multiome protocol to enable the fixation and permeabilization conditions required by Phospho-seq will not generate high-quality trimodal datasets.

We therefore pursued a computational alternative, integrating Phospho-seq and scRNA-seq datasets using our recently introduced ‘bridge integration’ procedure (Fig 3A) ^22^. We have previously shown that this workflow can successfully integrate distinct modalities collected in different experiments by leveraging a separately obtained multi-omic dataset as a ‘bridge’, even if the bridge dataset has reduced technical quality. The bridge integration procedure can successfully integrate data for both discrete cell types as well as continuous developmental trajectories, but requires that the multi-omic dataset is biologically representative (i.e. inclusive of all cell types and states) of the single-modality datasets. We emphasize that in our study, our bridge dataset was generated from the same set of samples, thereby satisfying this assumption and enabling us to integrate any future Phospho-seq datasets (i.e. with additional protein markers) without having to generate additional multi-omic data.

**Figure 3:**
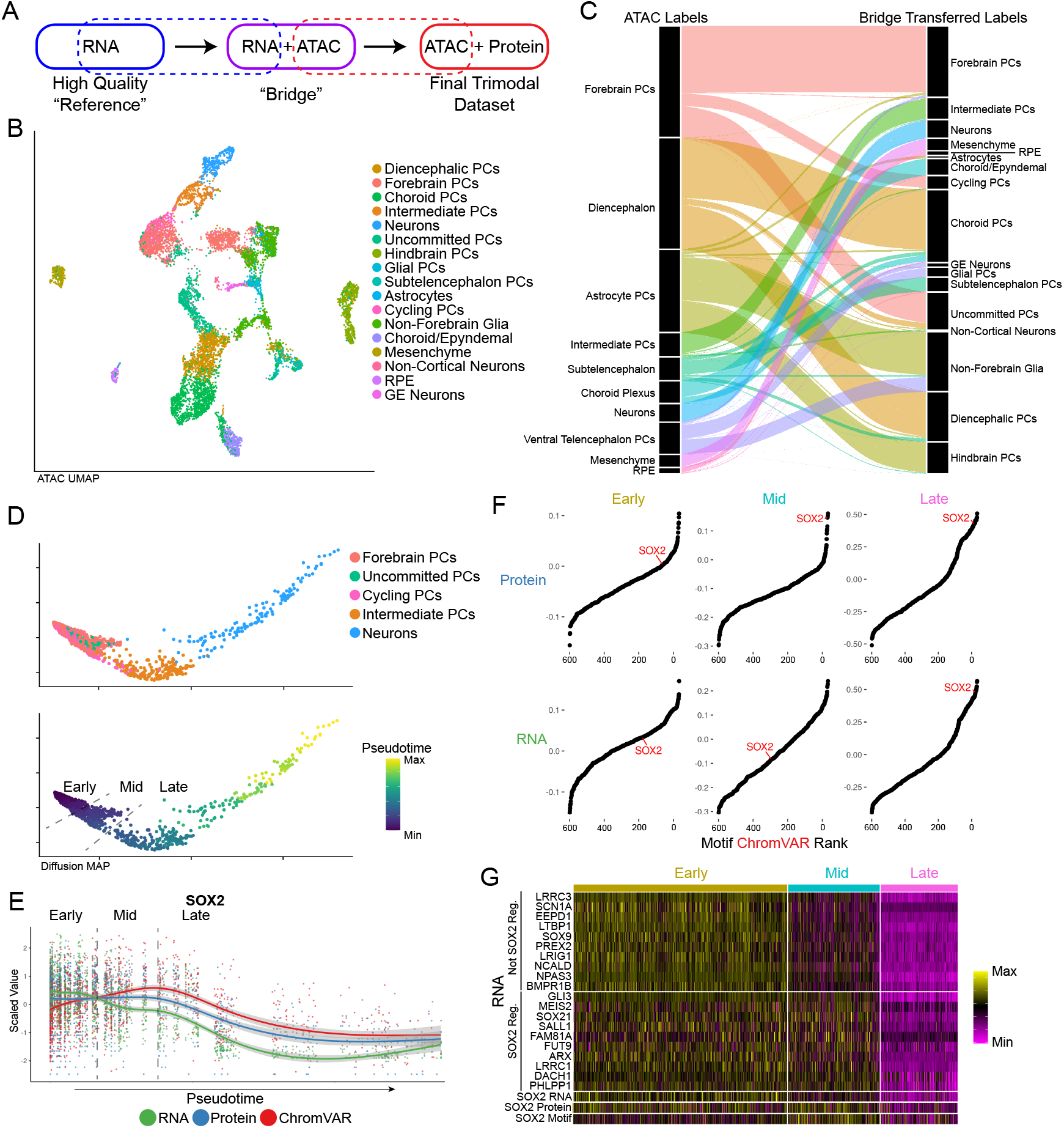
Bridge integration. **a)** Schematic of bridge integration. **b)** UMAP representation of cells based on ATAC-seq modality with cell type assignments from bridge integration. **c)** Alluvial plot demonstrating cell label transfer when using bridge integration. **d)** Diffusion map of cells differentiating from Forebrain PCs to Neurons colored by cell type (top panel) and pseudotime as determined by monocle (bottom panel). Dashed lines indicate pseudotime cut-offs determined by SOX2 expression and activity. **e)** Scatter plot showing scaled values of SOX2 RNA, protein and motif chromVAR score across pseudotime as determined in **(d)**. Dashed lines indicate the same SOX2-based pseudotime cut-offs as in **(d). f)** Rank-correlation plots of the correlation of transcription factor motif accessibility vs. SOX2 Protein (top panel) and SOX2 RNA (bottom panel) for each of the pseudotime cut-off group. **g)** Heatmap of a subset of genes not regulated by SOX2 and regulated by SOX2 across pseudotime bins.

Applying this workflow, we first identified 27 cell clusters from the scRNA-seq data and performed high-resolution manual annotation using canonical gene expression markers (Fig S3A-C). We next used bridge integration to annotate each Phospho-seq profile based on these labels (Fig 3B). We found that these transferred annotations were fully consistent with those originally derived from gene activity scores, but increased the granularity of annotation and interpretability. For example, Phospho-seq cells that were annotated as ‘Diencephalon’ fell into three categories after bridge integration: “Diencephalic PCs (Progenitor Cells), Choroid PCs, and Uncommitted PCs” (Fig 3C). After we performed bridge integration, we retrospectively identified differences in the chromatin accessibility profiles across high-resolution subsets that supported our transferred annotations. For instance, there were previously unobserved accessibility differences at canonical markers between progenitors and differentiated cells in both the choroid and astrocyte lineages (Fig S3D). Moreover, we obtained high prediction scores (70% > 0.75) for all cell types with the exception of astrocytes, which are one of the most rare cell populations in this system and were not sufficiently represented in the bridge data (Fig S3E). We therefore utilized our integrated manifold to ‘impute’ transcriptome-wide expression profiles for each Phospho-seq cell. We confirmed that these imputed profiles maintained high rank-correlation for TF motif accessibility, as well as measured ADT levels for the same gene (Fig S3F,G). These results confirm the applicability of bridge integration to obtain a high-quality RNA modality with Phospho-seq.

We leveraged our integrated dataset to explore the relationship between chromatin accessibility, gene transcription, and protein levels across a developmental trajectory of forebrain development. We subset our integrated Phospho-seq scRNA-seq dataset to include 1,578 cells that spanned a continuum of cell states from progenitor cells to neurons, and constructed a developmental trajectory based on diffusion maps (Fig 3D) ^35,36^. While SOX2 gene expression sharply decreased at initial stages of differentiation, we observed a developmental ‘lag’ in the decrease of downstream modalities including SOX2 ADT levels, and SOX2 transcription factor activity as estimated by chromVAR (Fig 3E). We identified a specific stage of the developmental trajectory where protein and RNA levels were discordant, and confirmed that in these cells, only SOX2 protein levels (and not RNA levels) were correlated with chromVAR scores (Fig 3F). While SOX2 is a well-established negative regulator of neuronal development ^37^, this analysis enabled us to identify genes whose downregulation preceded SOX2 protein downregulation, and are therefore unlikely to be direct or downstream targets of SOX2 itself, and vice-versa (Fig 3G and S3H,I). We conclude that integrated Phospho-seq data can perform multimodal characterization of the distinct temporal patterns and gene regulatory relationships that drive cellular dynamics.

### Using Phospho-seq data to identify gene regulatory networks in early neurodevelopment

By exploring relationships across modalities, multiomic datasets can help to reconstruct gene regulatory networks. While multiple groups have demonstrated how to leverage co-variation between chromatin accessibility and gene expression to link scATAC-seq peaks to the genes they regulate ^14,38,39^, our Phospho-seq data also allows us to link these peaks to transcription factors (TF) that regulate their accessibility. Recently, Argelaguet et al. ^40^ proposed ‘in-silco’ ChIP-seq, a computational approach to predict TF binding events from multiomic data, based on co-variation between the gene expression of a transcriptional regulator and an scATAC-seq peak. While this method assumes that RNA expression is a good proxy for protein levels, we reasoned that we could extend this analysis using protein measurements from our Phospho-seq panel.

For each TF, we performed a genome-wide search for peaks whose accessibility was highly correlated with TF protein abundance, and also harbored the TF binding motif (Fig 4A). As previously suggested ^40^, we used a high-resolution pseudo-bulking approach (‘SEAcells’) to reduce the sparsity of scATAC-seq data and identify more robust correlations ^41^ (Fig S4A). Furthermore, we identified a subset of these peaks whose accessibility was correlated (or anti-correlated) with the expression of the proximal gene, as inferred from our integrated dataset. This procedure enabled us to identify downstream activating and repressive targets for individual TFs, as well as specific enhancer elements that are likely to mediate these relationships.

**Figure 4:**
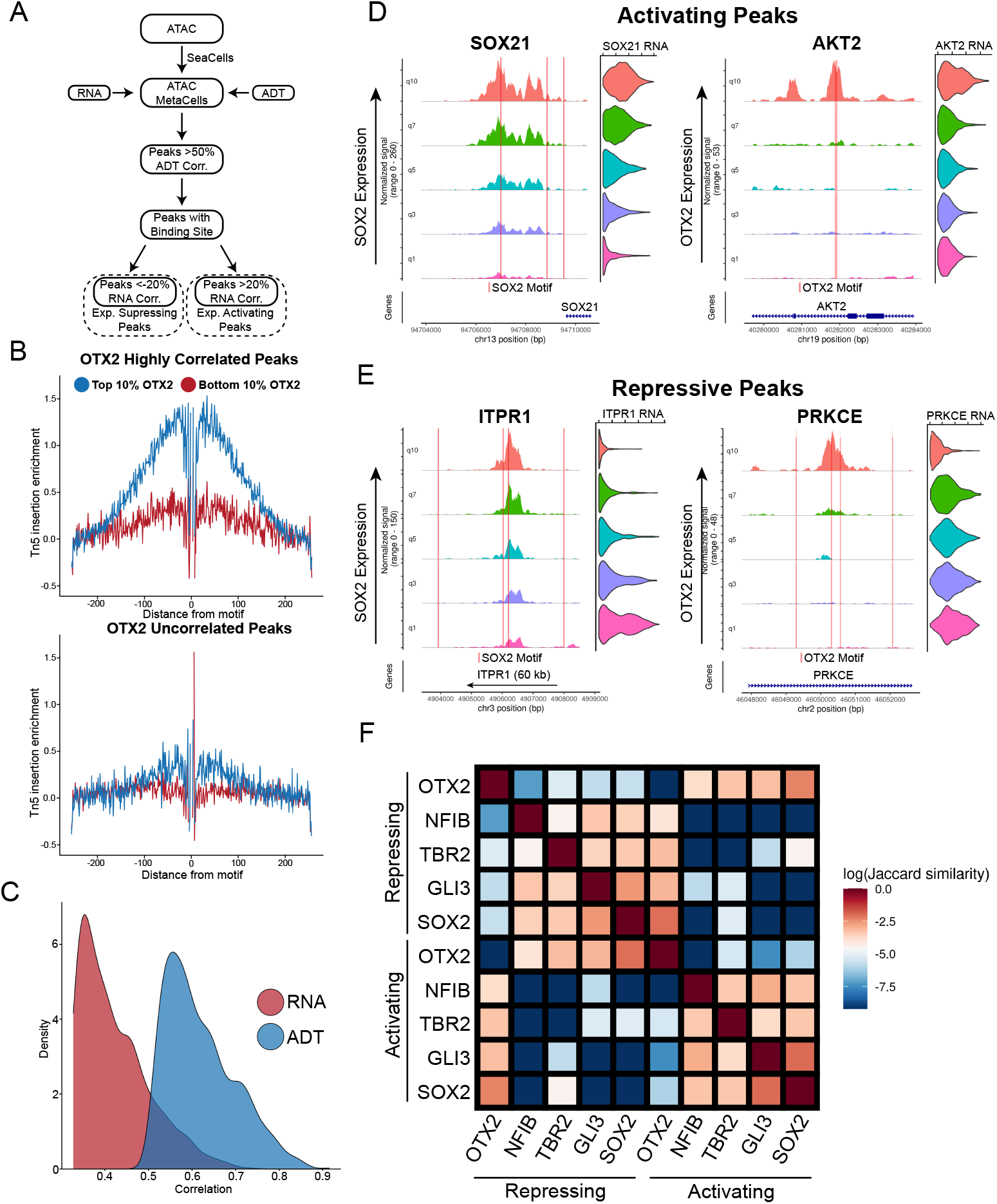
Transcription factor-specific cis-regulatory element discovery. **a)** Schematic of approach to use metacelling to discover cis-regulatory elements associated with individual proteins. **b)** Tn5 cut-site footprinting between cells with high OTX2 expression and low OTX2 expression in peaks that are highly correlated with OTX2 (top panel) and uncorrelated with OTX2 (bottom panel). **c)** Density of the correlations of top 1000 SOX2 correlated peaks when using SOX2 RNA from the multiome vs. SOX2 ADT. **d)** Examples of inferred transcription activating peaks associated with indicated proximal gene expression for SOX2 (left panel) and OTX2 (right panel). Coverage plots and violin plots are ordered by quantile ADT expression for each protein. Red lines indicated the location of a binding motif for each respective protein. **e)** Examples of inferred transcription repressing peaks associated with indicated proximal gene expression for SOX2 (left panel) and OTX2 (right panel). Coverage plots and violin plots are ordered by quantile ADT expression for each protein. **f)** Jaccard similarity matrix of inferred activating and repressing peaks from each of the TFs highlighted

We explored the performance of this procedure using OTX2 and SOX2, two transcription factors that are well-established regulators of neurodevelopment, and identified 5,033 and 2,562 linked peaks respectively. We first performed footprinting analysis, which aggregates the accessibility profiles across a collection of peaks. When performing this analysis in cells with high expression of the TF we observed a relative depletion of accessibility centered exactly at the TF binding site, which is strongly indicative of TF binding (Fig 4B and S4B,C). We did not observe evidence of binding when repeating this analysis in cells that did not exhibit TF protein expression, or when considering peaks whose accessibility was uncorrelated with TF levels. Moreover, when considering a previously published bulk ChIP-seq dataset ^42^ profiling SOX2 binding in *in vitro* differentiated human neural progenitors (hNPCs) and hESCs, we observed a clear elevation in SOX2 binding levels in our candidate peak set in the most analogous celltype, Forebrain PCs (Fig S4D). Finally, we observed that ADT-peak correlations for *bona fide* SOX2 peaks were substantially higher than RNA-peak correlations, reflecting both the higher quality and tighter biological associations with chromatin accessibility for protein measurements (Fig 4C and S4E).

Linking our candidate peaks to genes, we identified 1994 candidate targets (1287 activating, 707 repressive) for SOX2, and 3200 candidate targets (1772 activating, 1428 repressive) for OTX2 (Fig 4D,E; Table S2). Interestingly, we observed numerous cases where individual genes were regulated by multiple CREs, which were associated with different TFs. This included cases where putatively activating and repressive peaks were adjacently located within the same locus. We extended our analyses to consider additional TFs (TBR2, NFIB and GLI3; Fig S5A-F), and quantified the overlap between activated and repressed gene lists for different factors. This revealed synergistic relationships between some pairs of TFs, including SOX2 and GLI3, as well as antagonistic relationships between others (OTX2 and SOX2)(Fig 4F and S4F-H).

We found that Gene Ontology (GO) enrichments for predicted target genes were consistent with each TF’s known functional properties, including an enrichment for ‘axon guidance’ amongst targets of TBR2 (which is a positive regulator of late-stage neuronal differentiation ^43^), and an enrichment among SOX2 targets (radial glial cell regulator ^44^) for ‘glial cell differentiation’ genes (Fig S6A-J). For multiple TFs, we also observed an enrichment for genes associated with Wnt signaling, which is essential for brain patterning along the anterior-posterior axis. The pathway is known to increase activity in the posterior portion of the neural tube ^45,46^, supporting our observation of increased Wnt-related gene expression and increased chromVAR activities of WNT-responsive TFs in diencephalic cells compared to forebrain (Fig S6K,L). Lastly, we observed an enrichment for genes associated with ‘regulation of MAP kinase activity’ in OTX2 targets, which was supported not only by previous observations from mouse embryos where conditional deletion of OTX2 leads to decreases in phospho-MAPK/ERK signaling ^47^, but also by our Phospho-seq data which revealed a marked increase in phosphorylation of members of the MAPK/ERK cascade in OTX2+ PCs over OTX2-PCs (Fig 5A-D). We conclude that our integrated Phospho-seq dataset can effectively identify relationships across modalities and reconstruct gene regulatory relationships.

**Figure 5:**
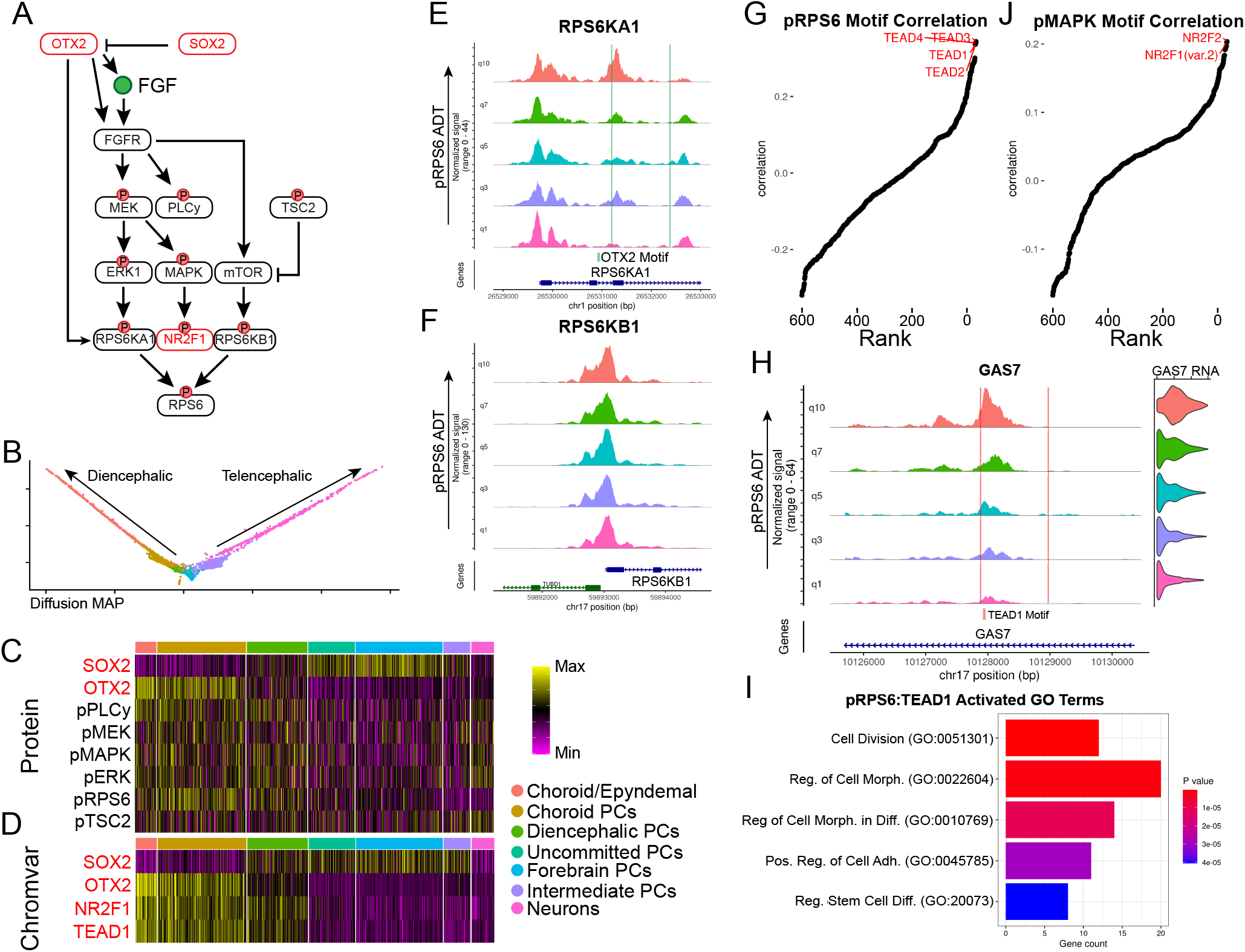
Differential signaling pathway activity across development. **a)** Schematic of SOX2 and OTX2 regulation of MAPK/ERK signaling. **b)** Diffusion MAP plot of diencephalic and telencephalic differentiation trajectories. Cell type colors correspond to legend in **(c). c)** Heatmap of ADT expression of proteins and phospho-proteins associated with MAPK/ERK and mTOR signaling. **d)** Heatmap of chromVAR scores for TF motifs enriched in diencephalic differentiation. **e)** Coverage plot of RPS6KA1 split into quantiles of pRPS6 levels across telencephalic and diencephalic differentiation trajectories with OTX2 binding motifs indicated. **f)** Coverage plot of RPS6KB1 split into quantiles of pRPS6 levels across telencephalic and diencephalic differentiation trajectories. **g)** Rank-correlation plot showing correlation between pRPS6 levels and motif accessibility across the whole Phospho-seq dataset. The top hits are indicated in red. **h)** Coverage and violin plots of GAS7 split by quantiles of pRPS6 levels across the whole dataset with TEAD1 motifs indicated by red lines. **i)** Bar plot of the top 5 most significant gene ontology categories associated with the top TEAD1 activated peak-gene links that are associated with pRPS6. **j)** Rank-correlation plot showing correlation between pMAPK levels and motif accessibility across the whole Phospho-seq dataset.

### Reconstructing signaling networks with Phospho-seq

While our initial analyses focused on TFs, our goal in developing Phospho-seq was to better characterize differences in signaling pathway activation across cell states, and to connect these changes to gene regulatory networks. For example, we found that pRPS6 was heterogeneously expressed across cell types. However, pRPS6 does not directly bind DNA, and instead works in concert with TFs to regulate gene expression through translation ^48^. Moreover, while pRPS6 is often utilized as an indicator for mTOR pathway activity, RPS6 can be phosphorylated by multiple kinases from different pathways - the ERK pathway via RPS6KA1 or the mTOR pathway via RPS6KB1 ^49^.

We found that our Phospho-seq data could help us better understand both upstream and downstream determinants of RPS6 phosphorylation. We found that the chromatin accessibility at the RPS6KA1 gene correlated with pRPS6 levels, as well as contained an OTX2 binding motif, while accessibility at RPS6KB1 was not correlated with pRPS6 levels (Fig 5E,F). While the kinase activity of these proteins are determined by upstream phosphorylation, the variability of pRPS6 levels appears to be in part driven by variation at the RPS6KA1 locus, indicating a more prominent role for ERK signaling over mTOR signaling in driving variable phosphorylation of pRPS6 in the diencephalic lineage.

When applying our ADT/motif rank correlation analysis, we identified a clear association between pRPS6 levels with multiple TEA/ATTS domain (TEAD) motifs (Fig 5G). TEAD transcription factors are the effectors of YAP-TAZ/Hippo signaling pathway, which is known to work in concert with MAPK/ERK signaling to regulate cell size ^50^, in cancer cells. Our findings link these pathways in neurodevelopment as well, and enabled us to identify 481 putative TEAD-regulated CREs (Fig 5H and Table S2). GO analysis of activating gene links revealed a strong enrichment for genes associated with cell division, growth, and morphogenesis, as would be expected for targets of both pathways (Fig 5I). Similarly, we found that pERK and pMAPK ADT levels were associated with the accessibility of NR2F1 and NR2F2 motifs (Fig 5J), representing an important neurodevelopmental regulator whose RNA expression was also highly elevated in cells with higher activation of MAPK/ERK signaling ^51,52^ (Fig S7A). These results again extend previously identified links between signaling pathways and TFs, originally identified in cancer cells, to cell state specification in neurodevelopment ^53^.

Finally, Phospho-seq can be used to measure phosphorylation among nuclear proteins, including transcription factors. Increased phosphorylation of the TF STAT3, which enhances its DNA-binding activity, is a characteristic feature of astrogliogenesis ^54^. In our dataset, we confirmed a high rank correlation of pSTAT3 with STAT3 motif accessibility, performing better than just STAT3 Protein or RNA alone (Fig S7B-D). We also saw an increase in pSTAT3 levels and STAT3 motif accessibility amongst glial cells in our dataset compared to progenitors or non-glial cells (Fig S7E,F), confirming previous biological observations ^55^. We then found 582 peak-gene links of CREs that harbored a STAT3 motif and whose proximal gene expression correlated with pSTAT3 levels. The expression of this gene set was specifically enriched in glial cells (Fig S7G), and included canonical astrocyte markers such as AQP4 (Table S2). We conclude that by exploring relationships across multiple molecular modalities, Phospho-seq can highlight the role of cell signaling pathways in neurodevelopmental fate specification.

## Discussion

Here we present Phospho-seq, a scalable approach to detect intracellular and intranuclear proteins, including phosphospecific states, alongside additional molecular modalities. We confirmed the sensitivity and specificity of our approach in cell lines, and subsequently applied Phospho-seq to profile three-month-old brain organoids. We added transcriptomic information to these data using ‘bridge integration’, allowing us to maximize the data quality obtained from each modality. With this integrated dataset we linked protein expression levels to the accessibility of *cis*-regulatory elements and the expression of proximal genes. We demonstrate how these connections can reconstruct gene regulatory relationships and reveal the causes and consequences of heterogeneous signaling during neurodevelopment.

We demonstrate that Phospho-seq is compatible with custom panels of user-conjugated antibodies, which can be easily and cost-effectively generated. While oligonucleotide-conjugated panels against cell surface proteins are readily available in both individual and pooled formats, these reagents are not widely available for intracellular targets. Moreover, Phospho-seq enables the user to perform multiplexed evaluation of the sensitivity and specificity for large panels of phospho-specific antibodies, by identifying correlations between ADT levels and chromatin/transcription states. We therefore expect that sequencing-based intracellular protein technologies will enable the identification and optimization of large intracellular panels, and that combining information across studies and biological systems will increase the generalizability of these results.

While pioneering technologies enable trimodal measurements of RNA, ATAC and surface protein or intranuclear abundance ^16,18,20^, in this study, we utilized ‘bridge integration’ ^22^ to harmonize molecular modalities collected in separate scRNA-seq and Phospho-seq datasets. Although this approach requires running additional experiments, we and others have found that simultaneous profiling of ATAC and RNA in cells or nuclei reduces the data quality associated with each modality ^38^. Improved fixation-compatible scATAC+scRNA methods may be helpful for addressing this issue in the future, but we note that a bridge-based experimental design may represent a flexible alternative for multimodal analysis where only a subset of samples need to be profiled with multiomic technologies. Importantly, we have previously shown that even small (<5,000 cell) 10x Multiome datasets can serve as effective bridges for large unimodal datasets, as long as they are biologically representative ^22^. Moreover, users of Phospho-seq may benefit from increasingly available reference atlases of either scRNA-seq or Multiome data ^56,57^, which would reduce the need for additional experimentation.

We anticipate that future studies will extend Phospho-seq to capture additional modalities related to chromatin state, and may shed additional light towards our understanding of how transcription factors regulate cellular chromatin. For example, combining Phospho-seq with scCUT&Tag ^58^ or NTT-seq ^59^ would identify TFs whose abundance correlated not only with chromatin accessibility, but with the presence of either activating or repressive chromatin marks. Further extensions that enable guide RNA capture (i.e. Perturb-ATAC ^60^, Spear-ATAC ^61^) would enable multiplexed genetic screens to utilize Phospho-seq to perform massively parallel identification and characterization of signaling regulators. Finally, applying Phospho-seq in concert with spatially-resolved profiling technologies ^62^ may shed light on both intercellular and intracellular signaling networks. We hope that the broad applicability of Phospho-seq will facilitate its adoption in diverse biological contexts, including development, immunology, neuroscience, and cancer, to discover how cell signaling determines cellular behavior and fate.

## Data availability

Seurat and Signac are freely available as open-source software packages at:

https://github.com/satijalab/seurat

https://github.com/stuart-lab/signac

Phospho-seq datasets generated for this manuscript are available at:

https://zenodo.org/record/7754315

## Acknowledgements

The authors would like to thank all the members of the Satija Lab and Treutlein Labs for thoughtful discussions related to this work, and specifically Sophie Jansen in the Treutlein Lab. J.D.B. is a postdoctoral fellow of the Jane Coffin Childs Memorial Fund for Medical Research. This work was supported by the Chan Zuckerberg Initiative (EOSS-0000000082, HCA-A-1704-01895 to R.S.), and the NIH (RM1HG011014-02, 1OT2OD026673-01, DP2HG009623-01, R01HD096770, R35NS097404 to R.S). B.T. was supported by the European Research Council (758877-Organomics), the Swiss National Science Foundation (Project Grant-310030_192604), F.Z. was supported by EMBO Long-Term Fellowship ALTF 36-2021.

---

**Figure S1.**
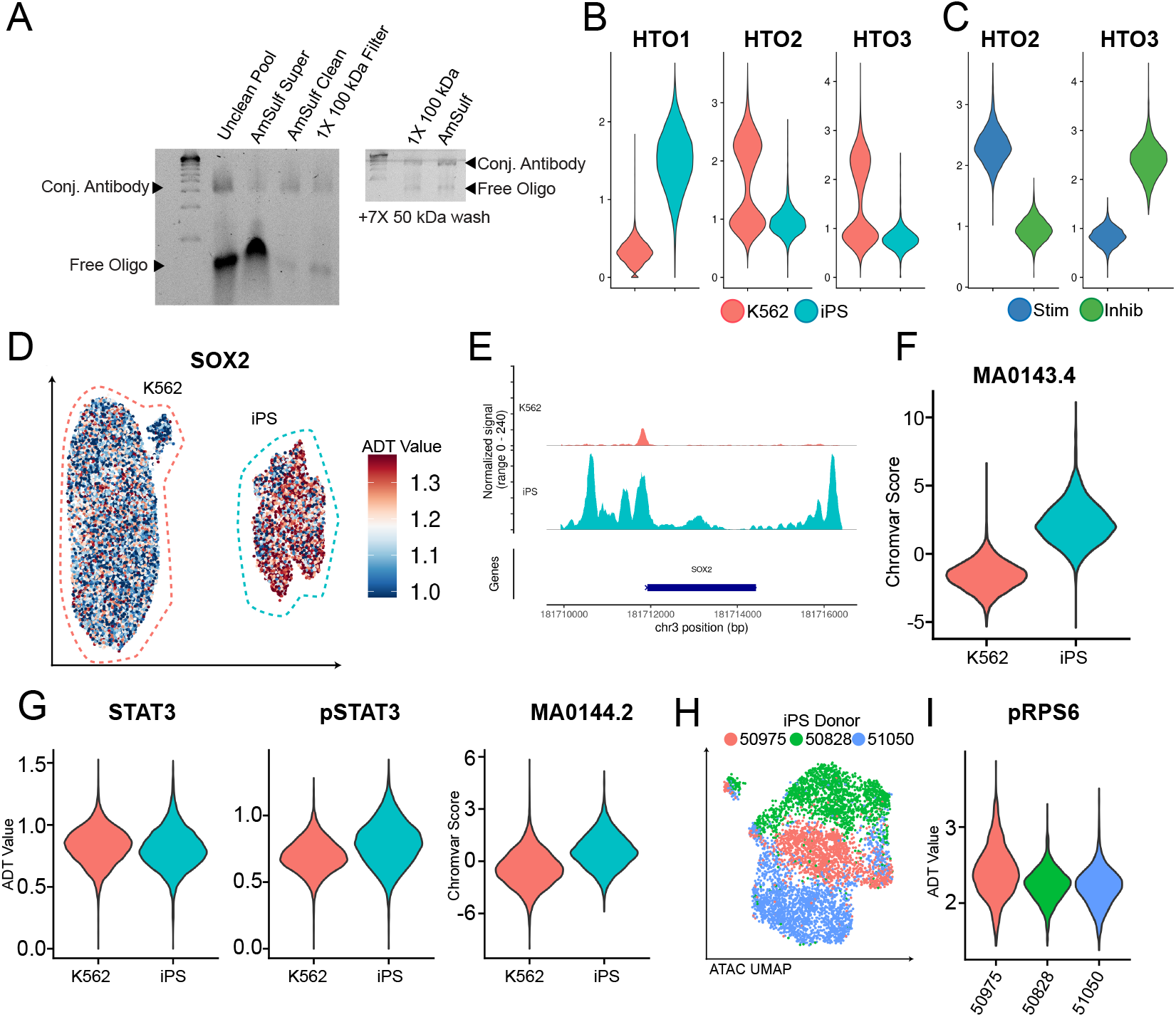
**a)** DNA electrophoresis gel of antibody pool at different indicated stages of clean-up showing the degree of remaining free oligo after filtration steps. **b)** Violin plot of HTO expression for each HTO in K562 and iPS cells. **c)** Violin plot of HTO expression for each HTO in K562 cells classified from stimulated or inhibited conditions. **d)** UMAP representation of K562 and iPS cells colored by normalized ADT values for SOX2. **e)** Coverage plot of chromatin accessibility of K562 and iPS cells at the SOX2 genomic locus. **f)** Violin plot of chromVAR scores for the SOX2 binding motif (MA0143.4) in K562 and iPS cells. **g)** Violin plots of normalized ADT values for STAT3 (left panel) pSTAT3 (middle panel) and STAT3 motif chromVAR scores (right panel) for K562 and iPS cells. **h)** UMAP representation of iPS cells colored by iPSC donor based on scATAC-sequencing data. **i)** Violin plot of normalized ADT values of pRPS6 split by iPSC donor identity.

**Figure S2.**
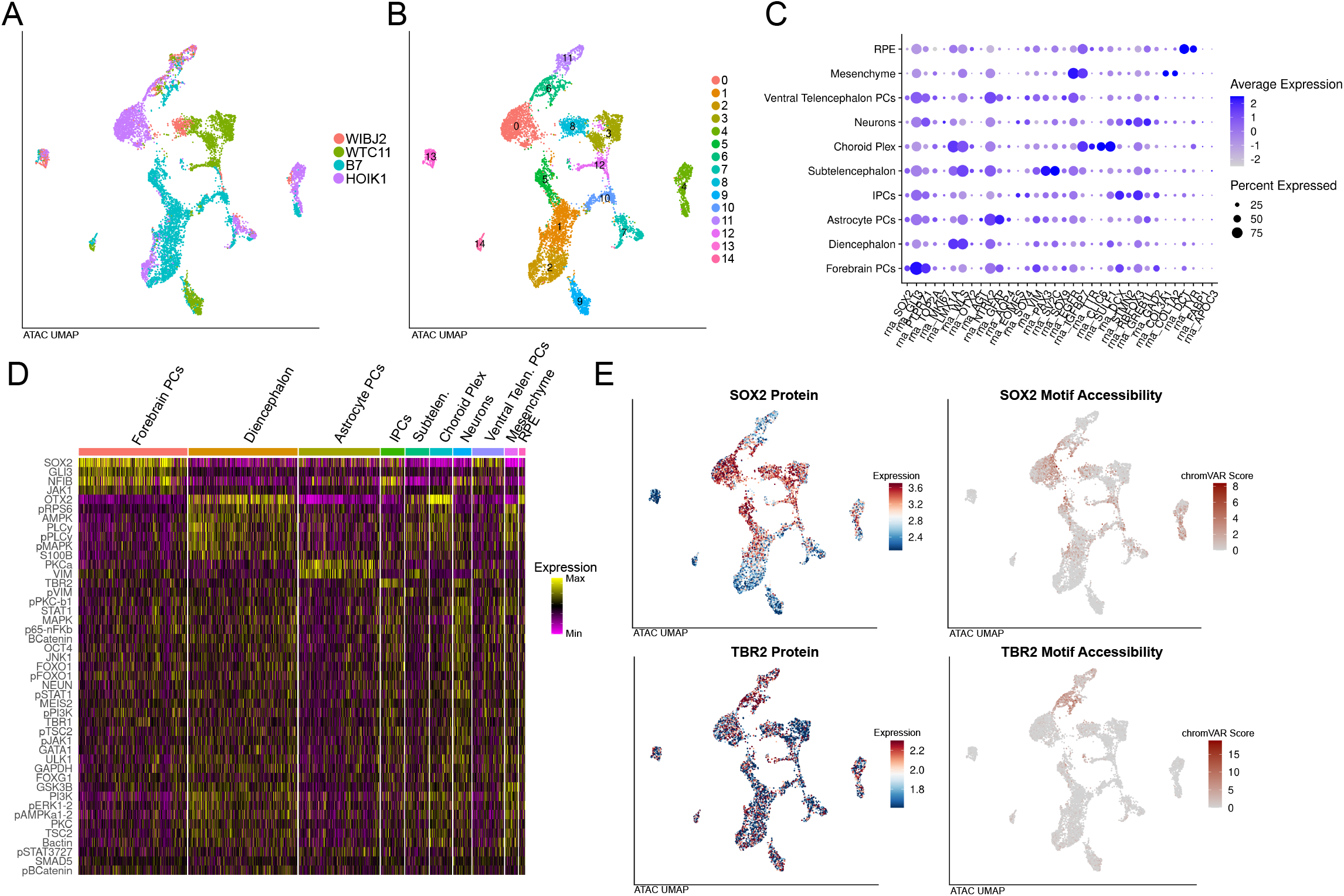
**a)** UMAP representation of cells colored by iPSC donor based on scATAC-Seq modality in Phospho-seq. **b)** UMAP representation of cells colored by unsupervised clustering based on scATAC-Seq modality in Phospho-seq. **c)** Dotplot showing the gene activity scores of marker genes and the percentage of cells they are expressed in from each assigned cell type identity in the Phospho-seq dataset. **d)** Heatmap showing scaled ADT expression for all differentially expressed ADTs organized by assigned cell type. **e)** UMAP representation of cells colored by normalized ADT values for SOX2 (upper left), chromVAR scores for SOX2 motif accessibility (upper right), normalized ADT values for TBR2 (lower left), chromVAR scores for TBR2 motif accessibility (lower right).

**Figure S3.**
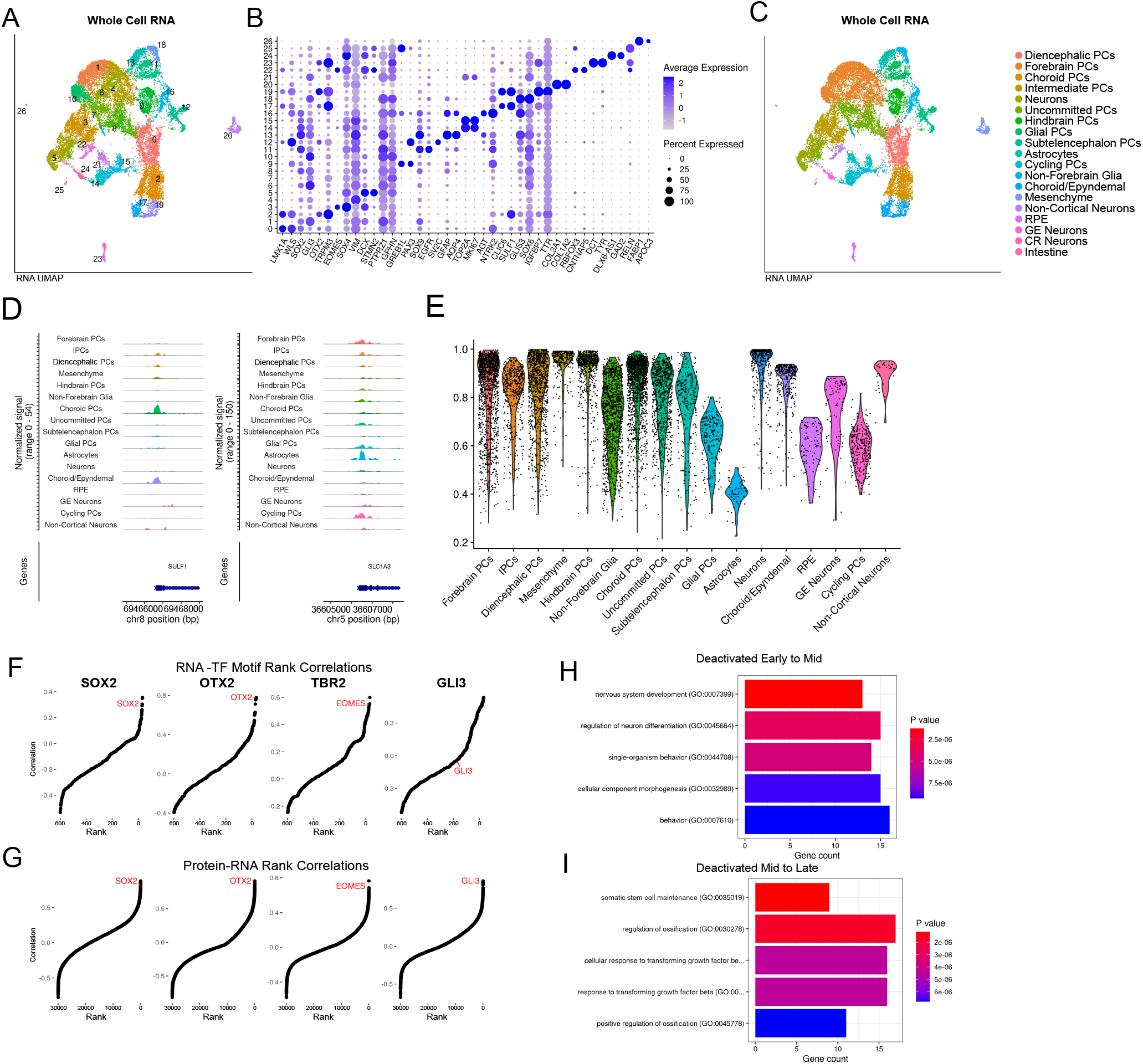
**a)** UMAP representation of cells colored by unsupervised cluster assignment from scRNA-Seq on whole unfixed cells. **b)** Dotplot showing the gene expression of marker genes and the percentage of cells they are expressed in from each cluster identity in whole, unfixed cells. **c)** UMAP representation of cells colored by cell type assignment from scRNA-Seq on whole unfixed cells. **d)** Coverage plots for SULF1 (left) and SLC1A3 (right) with transferred cell labels. **e)** Violin Plot of prediction scores per assigned cell type from bridge integration of whole cell RNA dataset with Phospho-seq dataset. **f)** Rank-correlation plots of imputed RNA vs TF-motif accessibility for each of the indicated genes. **g)** Rank-correlation plots of protein vs imputed RNA for each of the indicated proteins. **h)** Bar plot of the five most significant gene ontology categories associated with the top 500 genes decreasing from early to mid pseudotime bins across forebrain neuronal development. **i)** Bar plot of the five most significant gene ontology categories associated with the top 500 genes decreasing from mid to late pseudotime bins across forebrain neuronal development.

**Figure S4.**
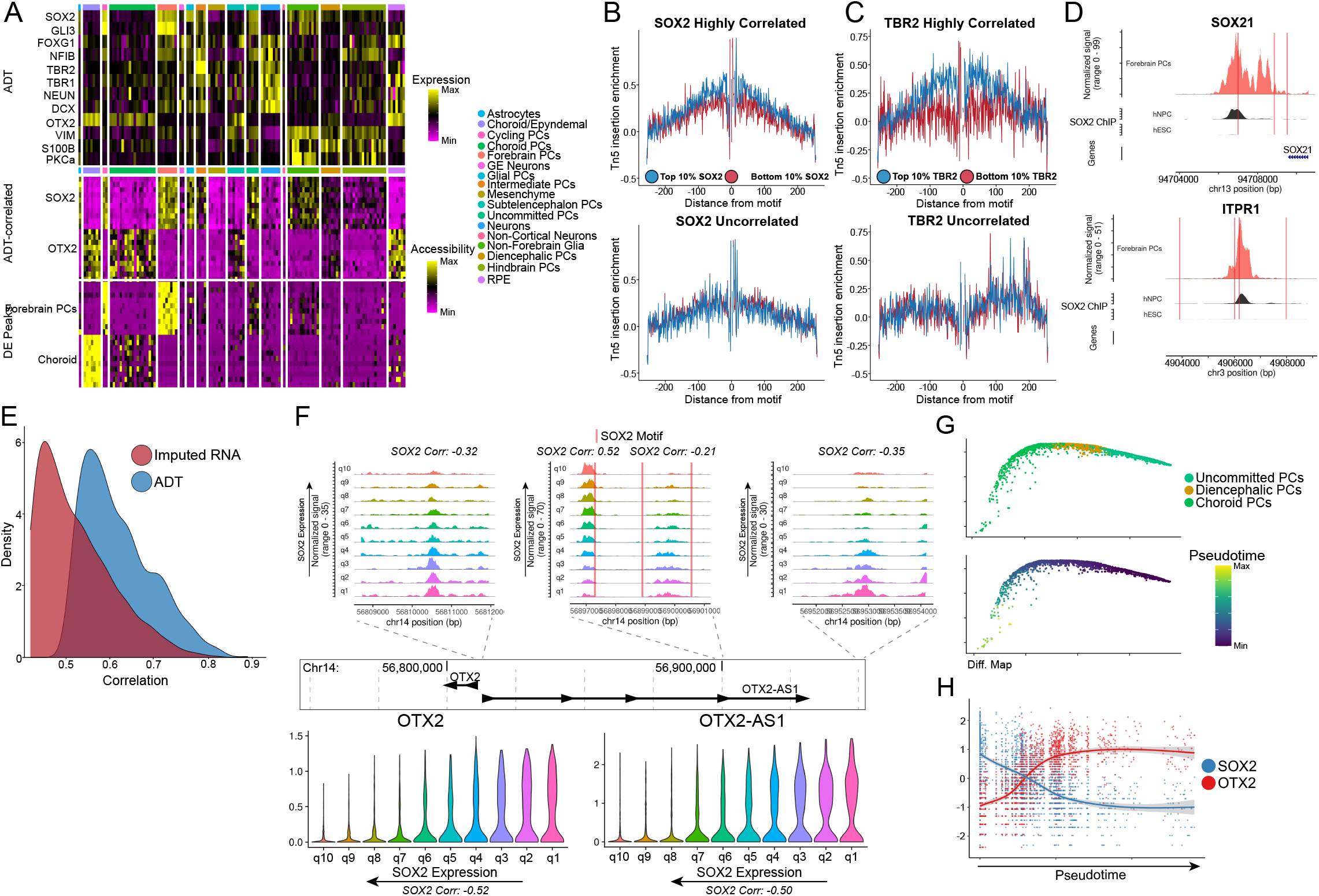
**a)** Heatmaps of metacelled data showing normalized marker ADT values (top panel), peaks correlated with SOX2 and OTX2 ADT expression (middle panel) and differentially accessible peaks for Forebrain and Choroid PCs determined from the non-metacelled data. **b)** Tn5 cut-site footprinting between cells with high SOX2 expression and low SOX2 expression in peaks that are highly correlated with SOX2 (top panel) and uncorrelated with SOX2 (bottom panel). **c)** Tn5 cut-site footprinting between cells with high TBR2 expression and low TBR2 expression in peaks that are highly correlated with TBR2 (top panel) and uncorrelated with TBR2 (bottom panel). **d)** Coverage plots of Forebrain PCs compared to peaks from a publically SOX2 ChIP-seq dataset from human neural precursors and human hESCs. **e)** Density of the correlations of top 1000 SOX2 correlated peaks when using SOX2 imputed RNA vs. SOX2 ADT. **f)** Combined plot of accessibility in peaks proximal to the OTX2 locus and associated gene expression. Plots are organized by quantile ADT expression of SOX2 throughout the whole dataset. Red lines on coverage plots are indicative of SOX2 binding motifs. **g)** Diffusion map of cells differentiating from Uncommitted PCs to choroid PCs colored by cell type (top panel) and pseudotime as determined by monocle (bottom panel). **h)** Scatter plot showing scaled values of SOX2 and OTX2 Protein across pseudotime as determined in **(g)**.

**Figure S5.**
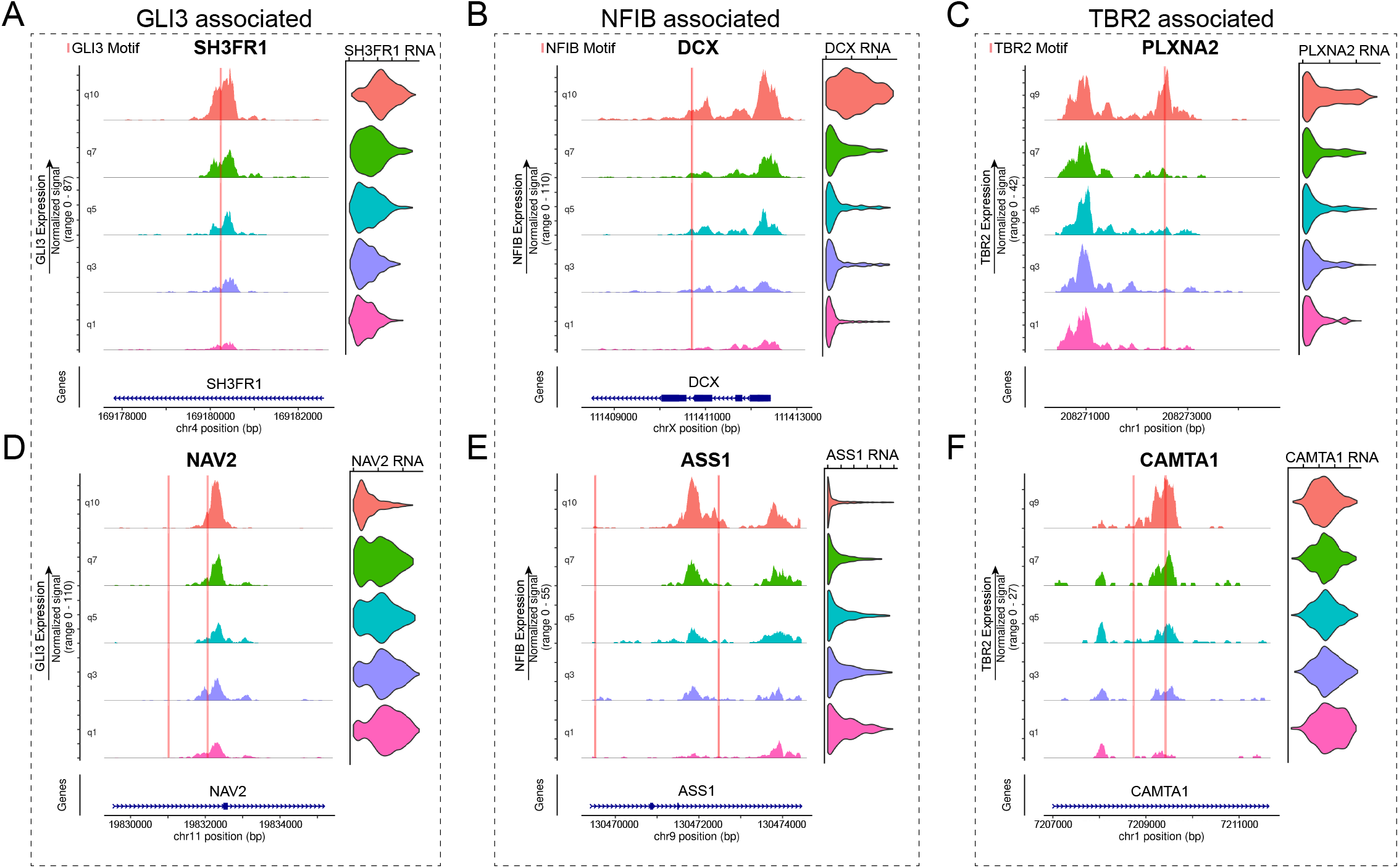
**a)** Example of an inferred transcription activating peak associated with GLI3. Coverage plots and violin plots are ordered by quantile ADT expression for GLI3. Red lines indicate the location of a GLI3 binding motif. **b)** Same as in **(a)** but for NFIB. **c)** same as in **(a)** but for TBR2. **d)** Example of an inferred transcription repression peak associated with GLI3. Coverage plots and violin plots are ordered by quantile ADT expression for GLI3. Red lines indicated the location of a GLI3 binding motif. **e)** Same as in **(d)** but for NFIB. **f)** same as in **(d)** but for TBR2.

**Figure S6.**
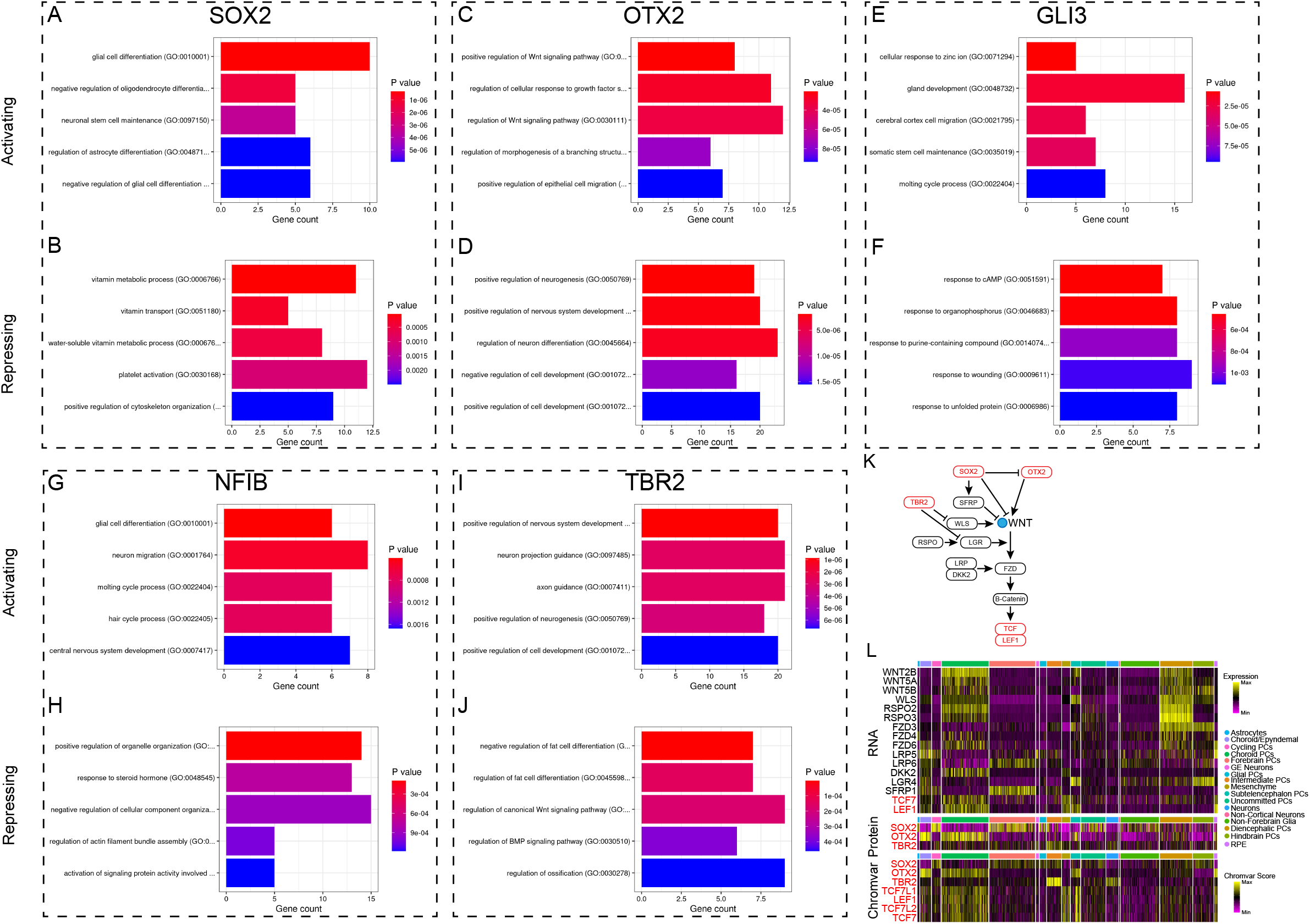
**a)** Bar plot of the top 5 most significant gene ontology categories associated with the genes in the top 500 peak-gene links that are SOX2 activated or repressed **(b). c)** Bar plot of the top 5 most significant gene ontology categories associated with the genes in the top 500 peak-gene links that are OTX2 activated or repressed **(d). e)** Bar plot of the top 5 most significant gene ontology categories associated with the genes in the top 500 peak-gene links that are GLI3 activated or repressed **(f). g)** Bar plot of the top 5 most significant gene ontology categories associated with the genes in the top 500 peak-gene links that are NFIB activated or repressed **(h). i)** Bar plot of the top 5 most significant gene ontology categories associated with the genes in the top 500 peak-gene links that are TBR2 activated or repressed **(j). k)** Schematic of SOX2,OTX2 and TBR2 regulation of WNT signaling. **l)** Heatmap of WNT-signaling related gene expression (top panel), SOX2,OTX2 and TBR2 protein expression (middle panel) and chromVAR scores for WNT-signaling related transcription factors

**Figure S7.**
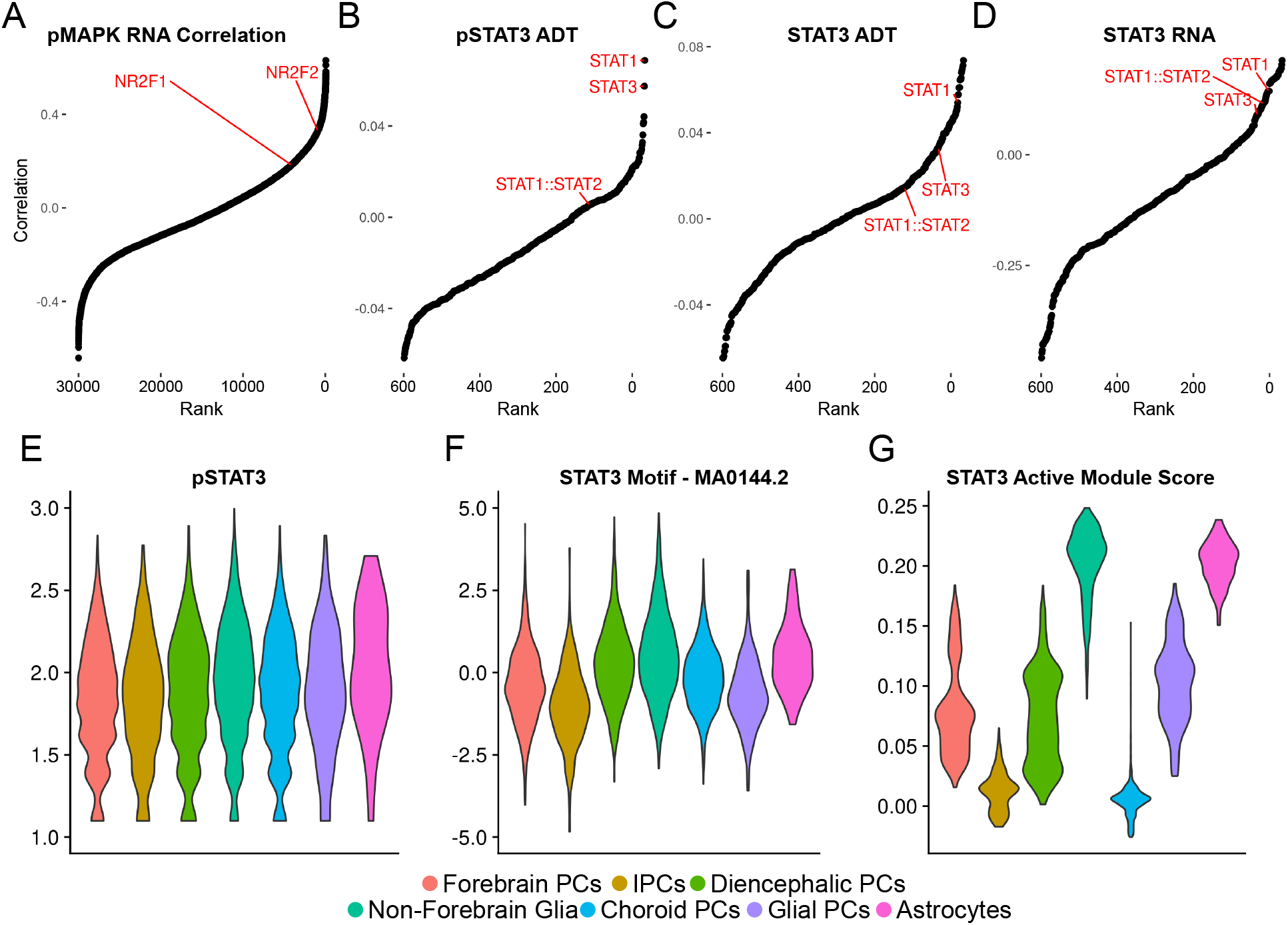
**a)** Rank-correlation plot of pMAPK ADT signal vs. RNA expression with NR2F1 and NR2F2 highlighted **b)** Rank-correlation plot of pSTAT3 ADT signal vs. TF motif accessibility with the STAT1 and STAT3 motifs highlighted. **c)** Same as in **(a)** but STAT3 ADT vs. TF motif accessibility. **d)** Same as in **(a)** but STAT3 RNA vs. TF motif accessibility. **e)** Violin plot of normalized pSTAT3 ADT levels in a subset of progenitor and glial cells. **f)** Violin Plot of STAT3 motif chromVAR scores in a subset of progenitor and glial cells. **g)** Violin plot of STAT3 activated gene module scores in a subset of progenitor and glial cells.

## References

1. Valls, P. O. & Esposito, A. Signalling dynamics, cell decisions, and homeostatic control in health and disease. Curr. Opin. Cell Biol. 75, 102066 (2022).

2. Badeaux, A. I. & Shi, Y. Emerging roles for chromatin as a signal integration and storage platform. Nat. Rev. Mol. Cell Biol. 14, 211–224 (2013).

3. Karin, M. & Smeal, T. Control of transcription factors by signal transduction pathways: the beginning of the end. Trends Biochem. Sci. 17, 418–422 (1992).

4. Thoreen, C. C. et al. A unifying model for mTORC1-mediated regulation of mRNA translation. Nature 485, 109–113 (2012).

5. Lee, M. J. & Yaffe, M. B. Protein Regulation in Signal Transduction. Cold Spring Harb. Perspect. Biol. 8, (2016).

6. Sturani, E. et al. Kinetics and regulation of the tyrosine phosphorylation of epidermal growth factor receptor in intact A431 cells. Mol. Cell. Biol. 8, 1345–1351 (1988).

7. Johnson, L. N. & Barford, D. The effects of phosphorylation on the structure and function of proteins. Annu. Rev. Biophys. Biomol. Struct. 22, 199–232 (1993).

8. Vandromme, M., Gauthier-Rouvière, C., Lamb, N. & Fernandez, A. Regulation of transcription factor localization: fine-tuning of gene expression. Trends Biochem. Sci. 21, 59–64 (1996).

9. Salazar, C. & Höfer, T. Multisite protein phosphorylation--from molecular mechanisms to kinetic models. FEBS J. 276, 3177–3198 (2009).

10. Blair, J. D., Hockemeyer, D. & Bateup, H. S. Genetically engineered human cortical spheroid models of tuberous sclerosis. Nat. Med. 24, 1568–1578 (2018).

11. Kumar, S. et al. Impaired neurodevelopmental pathways in autism spectrum disorder: a review of signaling mechanisms and crosstalk. J. Neurodev. Disord. 11, 10 (2019).

12. Krutzik, P. O. & Nolan, G. P. Intracellular phospho-protein staining techniques for flow cytometry: monitoring single cell signaling events. Cytometry A 55, 61–70 (2003).

13. Glassberg, J. et al. Application of phospho-CyTOF to characterize immune activation in patients with sickle cell disease in an ex vivo model of thrombosis. J. Immunol. Methods 453, 11–19 (2018).

14. Ma, S. et al. Chromatin Potential Identified by Shared Single-Cell Profiling of RNA and Chromatin. Cell 183, 1103–1116.e20 (2020).

15. Stoeckius, M. et al. Simultaneous epitope and transcriptome measurement in single cells. Nat. Methods 14, 865–868 (2017).

16. Chen, A. F. et al. NEAT-seq: simultaneous profiling of intra-nuclear proteins, chromatin accessibility and gene expression in single cells. Nat. Methods (2022) doi:10.1038/s41592-022-01461-y.

17. Rivello, F. et al. Single-cell intracellular epitope and transcript detection reveals signal transduction dynamics. Cell Rep Methods 1, 100070 (2021).

18. Mimitou, E. P. et al. Scalable, multimodal profiling of chromatin accessibility, gene expression and protein levels in single cells. Nat. Biotechnol. 39, 1246–1258 (2021).

19. Wong, M., Kosman, C., Takahashi, L. & Ramalingam, N. Simultaneous Quantification of Single-Cell Proteomes and Transcriptomes in Integrated Fluidic Circuits. Methods Mol. Biol. 2386, 219–261 (2022).

20. Swanson, E. et al. Simultaneous trimodal single-cell measurement of transcripts, epitopes, and chromatin accessibility using TEA-seq. Elife 10, (2021).

21. Chung, H. et al. Joint single-cell measurements of nuclear proteins and RNA in vivo. Nature Methods vol. 18 1204–1212 Preprint at https://doi.org/10.1038/s41592-021-01278-1 (2021).

22. Hao, Y. et al. Dictionary learning for integrative, multimodal, and scalable single-cell analysis. Preprint at https://doi.org/10.1101/2022.02.24.481684.

23. van Buggenum, J. A. G. L. et al. A covalent and cleavable antibody-DNA conjugation strategy for sensitive protein detection via immuno-PCR. Sci. Rep. 6, 22675 (2016).

24. Hao, Y. et al. Integrated analysis of multimodal single-cell data. Cell 184, 3573–3587.e29 (2021).

25. Fishman, J. B. & Berg, E. A. Ammonium Sulfate Fractionation of Antibodies. Cold Spring Harbor Protocols vol. 2018 db.prot099119 Preprint at https://doi.org/10.1101/pdb.prot099119 (2018).

26. Schep, A. N., Wu, B., Buenrostro, J. D. & Greenleaf, W. J. chromVAR: inferring transcription-factor-associated accessibility from single-cell epigenomic data. Nat. Methods 14, 975–978 (2017).

27. Bar, S. & Benvenisty, N. Epigenetic aberrations in human pluripotent stem cells. EMBO J. 38, (2019).

28. Kanton, S. et al. Organoid single-cell genomic atlas uncovers human-specific features of brain development. Nature 574, 418–422 (2019).

29. Fleck, J. S. et al. Inferring and perturbing cell fate regulomes in human brain organoids. Nature (2022) doi:10.1038/s41586-022-05279-8.

30. Yoon, S.-J. et al. Reliability of human cortical organoid generation. Nat. Methods 16, 75–78 (2019).

31. Lancaster, M. A. et al. Cerebral organoids model human brain development and microcephaly. Nature 501, 373–379 (2013).

32. Stoeckius, M. et al. Cell Hashing with barcoded antibodies enables multiplexing and doublet detection for single cell genomics. Genome Biol. 19, 224 (2018).

33. Huang, Y., McCarthy, D. J. & Stegle, O. Vireo: Bayesian demultiplexing of pooled single-cell RNA-seq data without genotype reference. Genome Biol. 20, 273 (2019).

34. Stuart, T. et al. Comprehensive Integration of Single-Cell Data. Cell 177, 1888–1902.e21 (2019).

35. Cao, J. et al. The single-cell transcriptional landscape of mammalian organogenesis. Nature 566, 496–502 (2019).

36. Angerer, P. et al. destiny: diffusion maps for large-scale single-cell data in R. Bioinformatics vol. 32 1241–1243 Preprint at https://doi.org/10.1093/bioinformatics/btv715 (2016).

37. Graham, V., Khudyakov, J., Ellis, P. & Pevny, L. SOX2 functions to maintain neural progenitor identity. Neuron 39, 749–765 (2003).

38. Trevino, A. E. et al. Chromatin and gene-regulatory dynamics of the developing human cerebral cortex at single-cell resolution. Cell 184, 5053–5069.e23 (2021).

39. Kartha, V. K. et al. Functional inference of gene regulation using single-cell multi-omics. Cell Genom 2, (2022).

40. Argelaguet, R. et al. Decoding gene regulation in the mouse embryo using single-cell multi-omics. Preprint at https://doi.org/10.1101/2022.06.15.496239.

41. Persad, S. et al. SEACells: Inference of transcriptional and epigenomic cellular states from single-cell genomics data. Preprint at https://doi.org/10.1101/2022.04.02.486748.

42. Zhou, C. et al. Comprehensive profiling reveals mechanisms of SOX2-mediated cell fate specification in human ESCs and NPCs. Cell Res. 26, 171–189 (2016).

43. Arnold, S. J. et al. The T-box transcription factor Eomes/Tbr2 regulates neurogenesis in the cortical subventricular zone. Genes Dev. 22, 2479–2484 (2008).

44. Pollen, A. A. et al. Molecular identity of human outer radial glia during cortical development. Cell 163, 55–67 (2015).

45. Polevoy, H. et al. New roles for Wnt and BMP signaling in neural anteroposterior patterning. EMBO Rep. 20, (2019).

46. Metzis, V. et al. Nervous System Regionalization Entails Axial Allocation before Neural Differentiation. Cell 175, 1105–1118.e17 (2018).

47. Mortensen, A. H., Schade, V., Lamonerie, T. & Camper, S. A. Deletion of OTX2 in neural ectoderm delays anterior pituitary development. Hum. Mol. Genet. 24, 939–953 (2015).

48. Bohlen, J., Roiuk, M. & Teleman, A. A. Phosphorylation of ribosomal protein S6 differentially affects mRNA translation based on ORF length. Nucleic Acids Res. 49, 13062–13074 (2021).

49. Roux, P. P. et al. RAS/ERK signaling promotes site-specific ribosomal protein S6 phosphorylation via RSK and stimulates cap-dependent translation. J. Biol. Chem. 282, 14056–14064 (2007).

50. Mugahid, D. et al. YAP regulates cell size and growth dynamics via non-cell autonomous mediators. Elife 9, (2020).

51. Faedo, A. et al. COUP-TFI coordinates cortical patterning, neurogenesis, and laminar fate and modulates MAPK/ERK, AKT, and beta-catenin signaling. Cereb. Cortex 18, 2117–2131 (2008).

52. Kanatani, S. et al. The COUP-TFII/Neuropilin-2 is a molecular switch steering diencephalon-derived GABAergic neurons in the developing mouse brain. Proc. Natl. Acad. Sci. U. S. A. 112, E4985–94 (2015).

53. Gay, F. et al. Multiple phosphorylation events control chicken ovalbumin upstream promoter transcription factor I orphan nuclear receptor activity. Mol. Endocrinol. 16, 1332–1351 (2002).

54. Moon, C. et al. Leukemia inhibitory factor inhibits neuronal terminal differentiation through STAT3 activation. Proc. Natl. Acad. Sci. U. S. A. 99, 9015–9020 (2002).

55. Bonni, A. et al. Regulation of gliogenesis in the central nervous system by the JAK-STAT signaling pathway. Science 278, 477–483 (1997).

56. Regev, A. et al. The Human Cell Atlas. Elife 6, (2017).

57. HuBMAP Consortium. The human body at cellular resolution: the NIH Human Biomolecular Atlas Program. Nature 574, 187–192 (2019).

58. Kaya-Okur, H. S. et al. CUT&Tag for efficient epigenomic profiling of small samples and single cells. Nat. Commun. 10, 1930 (2019).

59. Stuart, T. et al. Nanobody-tethered transposition enables multifactorial chromatin profiling at single-cell resolution. Nat. Biotechnol. (2022) doi:10.1038/s41587-022-01588-5.

60. Coupled Single-Cell CRISPR Screening and Epigenomic Profiling Reveals Causal Gene Regulatory Networks. Cell 176, 361–376.e17 (2019).

61. Pierce, S. E., Granja, J. M. & Greenleaf, W. J. High-throughput single-cell chromatin accessibility CRISPR screens enable unbiased identification of regulatory networks in cancer. Nat. Commun. 12, 1–8 (2021).

62. Deng, Y. et al. Spatial profiling of chromatin accessibility in mouse and human tissues. Nature 609, 375–383 (2022).

