## Supplementary Methods for "Phospho-seq: Integrated, multi-modal profiling of intracellular protein dynamics in single cells"

#### *Antibody Selection*

To select antibodies for use in the Phospho-seq assay, we used a hierarchical selection criteria based on available manufacturer's information. Antibodies were likely to be good candidates if they are first reported to work in intracellular flow cytometry and barring that, in immunocytochemistry on fixed cells. Antibodies that are reported to only work in western blot are less likely to work in Phospho-seq. Finally, the antibodies must be available in quantities of >5 μg in order to account for antibody loss in conjugation efficiency confirmation and post-conjugation processing. Antibodies from a variety of commercial vendors were used in this study including Abcam, Novus, R&D, ThermoFisher, Thomas Scientific and BioLegend. A full list of antibodies and catalog numbers is available in Table S1.

#### *Antibody Purification*

We used two methods of purifying antibodies from solutions with carrier proteins. For antibodies with BSA as a carrier protein, we used a BSA removal kit (Abcam: ab173231) following manufacturers instructions. For solutions that had more than just BSA contaminants (i.e. Ascites, gelatin), we used Magne Protein G beads (Promega: G7471). We followed manufacturer's instructions, eluting twice into a 100 mM Glycine-HCl solution (pH 2.7) and neutralizing with 2M Tris-buffer (pH 7.5).

#### *Antibody Conjugation and pooling*

For antibody conjugation steps prior to purification, we followed the protocol described here with minor alterations (https://citeseq.files.wordpress.com/2019/03/cite-seq_hyper_conjugation_190321.pdf), which leverages a cost-efficient and flexible conjugation strategy originally developed for immuno-PCR [1]. Briefly, TotalSeq-style oligos were ordered from IDT with sequences of /5AmMC12/CCTTGGCACCCGAGAATTCCAXXXXXXXXXXXXXXXBAAAAAAAAAAAAAAAAAAAAAAAAAAAA*A*A for hashing, and /5AmMC12/GTGACTGGAGTTCAGACGTGTGCTCTTCCGATCTNNNNNNNNNNXXXXXXXXXXXXXXXNNNNNNNNNGCTTTAAGGCCGGTCCTAGC*A*A for antibody tags where X's represent the antibody barcode and N's represent UMI tags. We attached a TCO-PEG4 linker to each oligo through incubation with TCO-PEG4-NHS reagent (Click Chemistry Tools: A137-25) and then column purified them with Micro Bio-Spin 6 Columns (Bio-Rad: 732-6221), confirming labeling through observation of a size shift compared to unlabeled oligos on a 4% agarose gel. The concentration of each oligo was quantified and diluted to 100 μM. As ordered, ~50 nmoles of oligo can be used for labeling ~3 mg of antibodies, thus requiring the labeling process to only be performed once per oligo-tag sequence for hundreds of experiments.

Next, 5 μg or more of purified antibody was labeled with mTz-PEG4 (Click Chemistry Tools: 1069-10) and conjugated to the TCO-PEG4 labeled oligo at a target concentration of 15 pmol per 1 μg of antibody. This leaves each antibody conjugated to 2-4 copies of each oligo on average. Each conjugation was confirmed through running 1 μg of conjugated antibody on a 4-12% protein gel (Thermo: XP04125BOX) and observing a laddered size shift in increments of 30 kDa above the unconjugated IgG size of 155 kDa. The final concentration of each conjugated antibody was estimated through a BCA assay (Thermo: 23227).

Finally, for each experiment, 3-5 μg of each antibody was pooled together and treated with 40% saturated (~4.32 M) Ammonium Sulfate to salt-out the conjugated antibodies from the unconjugated oligos still present in the antibody solution. After the majority of the remaining oligos were removed, the pooled solution was passed through a 50 kDa MWCO filter (EMD Millipore: UFC505024) 5-7 times - checking the flowthrough solution for ssDNA (leftover oligos) concentration after each centrifugation. When the flowthrough solution is at or below 0.2 ng/ul, the leftover antibody is resuspended from the filter and checked on a 4% agarose gel for two bands, one lower oligo band (90 nt) and one higher antibody band (~400 nt), with the aim of having the oligo band as faint as possible. The final antibody pool was then quantified through a BCA assay.

*Cell Culture*

K562 cells were maintained in IMDM media supplemented (ThermoFisher: 31980030) with 10% FBS (Corning:35-010-CV), 1 mM Pen-Strep (Sigma: P0781-50ML) and 1X Non-essential amino acids (Sigma:M7145-100ML). Four hours before harvesting, K562 cells were removed from maintenance media, centrifuged 5 minutes at 1000 rpm, and resuspended in IMDM media with no supplements. Cells were counted and $1x10^6$ cells were transferred into two separate flasks. In one flask, cells were immediately treated with 500 nM PX-866 (Cayman Chemical: 13645) while in the other flask, 50 ng/ml EGF (R&D Systems: 236-EG-200) was added 15 minutes before harvest.

Three lines of iPSCs were obtained from the New York Stem Cell Foundation ( 51050,50828,50975). Cells were thawed into feeder-free conditions with mTESR1 (STEMCELL: 85850) and 10% CloneR (STEMCELL: 05888) with a Geltrex substrate (Fisher: A1413301). After two days, cells were changed into maintenance media – mTESR1 supplemented with 1 mM Pen-Strep. Throughout culture, media was changed daily and cell colonies were passaged by treatment with 1:3 Accutase (STEMCELL: 07920) diluted with PBS (Caisson Labs:PBL06-500ML) for five minutes. Cell lines were passaged a maximum of five times before an experiment in order to reduce the number of cell-culture related chromosomal abnormalities.

*FACS*

iPS cells were harvested through dissociation with accutase for 5 minutes and centrifuged in growth media at 1000 rpm for 5 minutes. K562 cells in suspension were harvested through centrifugation in growth media. Both cell lines were resuspended in 1 ml PBS + 0.04% BSA (Fisher: BP9706-100) and run through a 40 μm filter to obtain a single cell suspension. These suspensions were further centrifuged for 5 mins at 300 x g in a tabletop swinging bucket centrifuge. Cells were resuspended in 450 μl of PBS and were fixed by adding 30 μl of 16% formaldehyde (Sigma: F8775-25ML) (final concentration, 1% FA in PBS). Cell suspensions were left to fix for 10 minutes at RT, with inversion every 3 minutes. The fixation reaction was quenched by adding 68.5 µl 1M glycine and filling the tubes with ice-cold PBS. The suspensions were centrifuged for 5 minutes at 400 x g at 4C, after which the supernatant was removed. The cells were resuspended in 1 ml PBS was added and the centrifugation was repeated. After the second centrifugation, the cells were resuspended in 100 ul of lysis buffer (10 mM Tris-HCl, 10 mM NaCl, 3.33 mM $MgCl_2$, 0.1% NP-40 (Thermo: 28324), 1% BSA in $H_2O$) and incubated on ice for 5 minutes for permeabilization. After 5 minutes, wash buffer #1 (10 mM Tris-HCl, 10 mM NaCl, 3.33 mM $MgCl_2$, 1% BSA in $H_2O$) was added up to 1 ml and cells were centrifuged for 5 minutes at 500 x g after which the supernatant was discarded and the cells were resuspended in staining buffer (3% BSA in PBS) for blocking and placed in a tube rotator at RT for 30 minutes.

For cells that received a primary antibody bound by EcoRI single-stranded DNA binding protein (SSB) (Promega:M3011), the SSB binding reaction happened during the blocking step. Briefly, 1 µg of antibody was incubated with 8 µg of SSB in a solution of 1x NEB buffer 4 (NEB:B7004S) for 30 mins at 37 C. After this incubation, BSA and PBS were added to make a final solution of 3% BSA in 1X PBS.

For staining, the blocking buffer was removed through centrifugation at 600 x g for 5 minutes and washing in wash buffer #2 (3% BSA in PBS + 0.1% Tween) three times. Cells were then resuspended in 100 µl of staining buffer with the appropriate antibodies and placed on the tube rotator for 1 hour at RT. After this 1 hour, the cells were washed three times in wash buffer #2 and then resuspended in staining buffer with a compatible secondary antibody (BioLegend: 406605,407507,406707,408205) and stained for 30 minutes in the dark. Following three more washes, the cells were resuspended in MACS buffer (2 mM EDTA and 0.5% BSA in PBS) and run through the flow cytometry protocol on a Sony MH800 cell sorter with secondary only stained K562 cells acting as the negative control and a goal of 20,000 cells per condition. Flow cytometry analysis and plotting was performed using FlowJo v9.3.2.

*Phospho-seq*

The fixation and permeabilization steps for the Phospho-seq experiments proceeded the same as in FACS up until the blocking step, at which point in Phospho-seq, we also performed cell hashing [2]. For the pilot experiment, the iPS, K562-Stim and K562-Inhib were separately stained with unique, home-conjugated hashing antibodies, while for the organoid experiments, the fixed permeabilized cells were split equally into four tubes for hashing. Briefly, 1 ug of TotalSeqA-conjugated CD298 and B2M antibody mix was added into the blocking buffer. Additionally, we included 100 µg of 30-nt 3' blocked single stranded DNA oligo to act as an additional block within the cells ( NNNNNNNNNNNNNNNNNNNNNNNNNNNNNN/3ddC/). For the organoid experiments we also included an additional 4 nM of four mixed unconjugated TSA oligos to act as a permeabilization control measure. where we could normalize cells with higher degrees of permeabilization and thus more potential for antibody inflow, against a common background of freely binding oligos [3]. Cells were blocked for 30 minutes at room temperature on a tube rotator. During this blocking step, the primary antibody pool was incubated with SSB as above in preparation for primary antibody staining. For the pilot experiments, we used 0.5 µg/antibody for primary staining while for the cerebral organoid experiment we used 0.25 µg/antibody. After blocking, cells were centrifuged at 600 x g for 5 minutes and washed once in wash buffer #2. Cells from each partition were resuspended in wash buffer #2, counted and combined in equal proportions up to 1 million cells total. Combined cells were centrifuged again at 600 x g for 5 minutes, resuspended in the primary antibody solution and placed in a tube rotator for 1 hour at RT. After primary antibody staining, cells were washed twice in wash buffer #2 and resuspended in 1x nuclei buffer (10X). The cell suspension was sent through a 40 µm filter and centrifuged at 600 x g for 5 minutes. At this point, enough supernatant was removed to leave ~50 µl cell suspension. 5 µl of this cell suspension was used for quantification and the concentration was adjusted to 6,000 cells/µl if possible. 5 µl of this solution (30,000 cells) was added to a solution of 7 µl ATAC buffer B and 3 µl ATAC enzyme (10X) and the rest of the protocol followed 10X scATAC kit v1.1 (10X Genomics: 100175) protocols with minor alterations in line with the ASAP-Seq protocol (below).

After tagmentation, 0.5 µl each of bridge oligo A and bridge oligo B was added to the barcoding reaction to allow for TotalSeq-A and TotalSeq-B oligo capture. To allow for better hybridization between the capture oligo, bridge oligo and antibody tags there was a 5 minute 40 C step added at the beginning of the GEM incubation. After silane bead elution (Step 3.1o), 43.5 µl instead of 40.5 µl Elution Solution I was added for elution - 40 µl was kept for regular SPRI cleanup while 3 µl was set aside for for added complexity in the tag library PCR. After the first SPRI cleanup (Step 3.2d), the supernatant was saved instead of discarding. This supernatant contains all of the captured hashing and antibody tags, while the DNA on the beads contains the ATAC fragments and can proceed normally as written. The supernatant was treated with 32 µl of SPRI beads, allowed to bind for 5 minutes and subsequently cleaned up with 80% EtOH and eluted in 42 µl Buffer EB. This elute was combined with the 3 µl set aside from above and split into two tubes for amplification of TotalSeq-A and TotalSeq-B respectively. These two PCRs commenced separately with conditions as follows: 50 µl 2x KAPA HotStart ReadyMix (KAPA biosystems:07958935001), 2.5 µl 10 µM P5 primer (AATGATACGGCGACCACCGA), 10 µM P7 primer ( CAAGCAGAAGACGGCATACGAGATXXXXXXXXGTGACTGGAGTTCCTTGGCACCCGAGAATTCCA for TotalSeq-A and CAAGCAGAAGACGGCATACGAGATXXXXXXXXGTGACTGGAGTTCAGACGTGTGC for TotalSeq-B, where Xs are i7 index nucleotides), 22.5 µl $H_2O$ and 22.5 µl input fragments. The thermal cycler program is: 95C for 3 minutes, 14-18 cycles of 95C - 20 seconds, 60C - 30 seconds, 72C - 20 seconds, then finally 72C for 5 minutes. These reactions were then cleaned up with 1.6X SPRI beads and eluted in 20 µl Buffer EB.

*Sequencing and Alignment*

Assembled sequencing libraries were quantified by a Bioanalyzer High-Sensitivity DNA chip (Agilent: 5067-4626) and combined at a molar proportion of 25% ADT, 10% HTO and 65% ATAC. These combined libraries were sequenced on a NextSeq 550 instrument using a 75-cycle high-output kit (Illumina: 20024906) with 34 bp Read 1, 8 bp i7, 16 bp i5 and 34 bp Read 2 parameters. ATAC Sequencing data was demultiplexed and aligned to the hg38 reference genome using CellRanger-ATAC v2.0.0 (10X genomics). ADT, HTO and Free TotalSeq-A tag sequencing data was demultiplexed and aligned to index references using alevin [4].

*Data Processing for Pilot Experiment*

Chromatin data was processed with Signac v1.9 [5]. We performed peak calling using the MACS3 algorithm [6], and performed downstream analysis using standard workflows (TFIDF, SVD and LSI for dimensionality reduction) and default parameters. ChromVAR [7] was called within Signac for TF accessibility. HTOs were normalized using CLR normalization, and we performed demultiplexing and doublet detection using the HTOdemux function with default parameters. Only cells classified as singlets were kept, leaving 14,931 cells. Additional demultiplexing to assign cells to individual iPS donors was performed using Vireo v0.5.6 [8] , and relied on variant calling from 1000 Genomes Project and GnomAD common variants using Vartrix v1.1.22. ADTs were processed using default parameters from Seurat, including CLR normalization.

*Organoid Culture and Harvesting*

We used four human iPS cell lines (Hoik1 and Wibj2 from the HipSci resource8, 01F49i-N-B7 (B7) from Institute of Molecular and Clinical Ophthalmology Basel, and WTC from the Allen Institute). Stem cell lines were cultured in mTESR1 and supplemented with penicillin–streptomycin (1:200, Gibco: 15140122) on Matrigel-coated plates (Corning: 354277). Cells were passaged 1–2 times per week after dissociation with TrypIE (Gibco: 12605010) or EDTA in DPBS (final concentration 0.5 mM) (Gibco:12605010). The medium was supplemented with Rho-associated protein kinase (ROCK) inhibitor Y-27632 (final concentration 5 µM, STEMCELL Technologies: 72302) the first day after passage. Cells were tested for mycoplasma infection regularly using PCR validation (Venor GeM Classic, Minerva Biolabs) and found to be negative. A total of 2,000–3,000 cells were aggregated in ultra low-attachment plates (Corning: CLS7007) to generate brain organoids using a whole-brain organoid differentiation protocol [9].

Single-cell suspensions were acquired by dissociation of the organoids with a papain based dissociation (Miltenyi Biotec: 130-092-628). The organoids were cut into pieces and washed with HBSS (no magnesium, no calcium, Gibco: 14170120 ) to remove debris before prewarmed papain solution (2 ml) was added and incubated for 15 min at 37 °C. Enzyme mix A was added before the tissue pieces were triturated 5–10 times with 1,000 µl wide-bore and P1000 pipette tips. The tissue pieces were incubated twice for 10 min at 37 °C with trituration steps in between and after with P200 and P1000 pipette tips. Cells were filtered consecutively with a 30 µm pre separation filter and centrifuged. Cells were counted and 1-3 million cells pre sample were cryopreserved in CryoStor CS10 (STEMCELL Technologies: 07952) at −80 C before moving to liquid nitrogen storage the following day.

On the day of the experiment, cells were removed from cryopreservation and thawed at 37C. Cells were deposited into Neurobasal media (Thermo: 21103049) supplemented with 10% FBS and centrifuged for 5 mins at 300 x g. Cell pellets were resuspended in 450 µl PBS, run through a 40 µm filter, after which fixative was added and the Phospho-seq experiment continued as written above.

*Initial Data Processing for Organoid Experiment*

Chromatin data was quantified, normalized, and processed using the same workflow as for the Phospho-seq pilot experiment. We removed one cluster of cells that was defined by low sequencing depth. We assigned each cell to its organoid line and donor-of-origin using Vireo v0.5.6[8] , and relied on variant calling from 1000 Genomes Project and GnomAD common variants using Vartrix v1.1.22.

For ADT processing, cells with over 10,000 ADT counts were removed from the analysis, as these were strong outliers in ADT data. This left 9,034 cells total in the final dataset. ADT signal was normalized using scTransform v2 [10] and regressed against the number of free TotalSeq-A counts, which represent the degree of cell permeabilization. To identify relationships between ADT levels and TF activity scores, we calculated Pearson correlations between each ADT value and each TF motif chromVAR scores across all cells. These correlations were then plotted according to their rank compared to every other TF motif in the JASPAR2020 motif dataset.

*Bridge Integration*

Additional single cell experiments were performed for bridge integration on the same Day 90 organoid samples. For the bridge dataset, nuclei were extracted from cells and run through the 10X Multiome ATAC + Gene Expression protocol according to manufacturer's instructions (10X Genomics:1000285). Libraries were sequenced on a NextSeq 550 instrument using a 150 cycle high-output kit with 50 bp R1, 8 bp i7, 16 bp i5 and 86 bp R2. Data was aligned using CellRanger-arc v 2.0.0 to hg38. Aligned data was processed using default parameters in Seurat and Signac including a cutoff of 500 counts and SCT normalization for the RNA modality. Nuclei were demultiplexed using variant calling on the ATAC modality and only singlet nuclei were used. This left 4,958 nuclei in the bridge dataset.

For the whole cell scRNA-Seq dataset, whole, unfixed cells were hashed with four homemade hashing antibodies, loaded at 25,000 cells/lane and run through the 10X scRNA-Seq v3.1 kit according to manufacturer's instructions (10X Genomics: 1000268). Separate HTO and RNA libraries were constructed and sequenced on a NextSeq 550 instrument at a ratio of 90% RNA and 10% HTOs using a 75-cycle high-throughput kit with 28 bp R1, 8 bp i7 and 56 bp R2. Transcriptomic data was aligned using CellRanger v7.0.0 to hg38 and HTO data was aligned to indices using Alevin. Cells were processed using default parameters in Seurat with SCT normalization. HTOs were quantified and cells were clustered according to HTO expression, with clusters representing doublets removed from subsequent analysis. This processing resulted in 19,280 cells in the final reference dataset.

Bridge integration and RNA imputation for the Phospho-seq dataset was performed using the standard bridge integration workflow [11], described here: https://satijalab.org/seurat/articles/bridge_integration_vignette.html. First we re-called peaks in the Phospho-seq ATAC dataset using only those peaks present in the bridge dataset. This procedure identifies 'anchors' between the Phospho-seq and scRNA-seq dataset, after representing them both based on the same multi-omic dictionary. After the transfer anchors were determined, the cell labels and RNA modality were transferred onto the Phospho-seq dataset using the TransferData function in Seurat.

*Pseudotime and trajectory analysis*

To look for gene expression, protein and chromVAR patterns across a trajectory, cells were first subset by both donor and cell type lineage - Hoik1 cells and forebrain neuronal lineage or B7 cells and diencephalonic lineage. The 'destiny' [12] package in R was used to create diffusion maps based on the lsi embeddings from these subset datasets. Monocle3 [13] was then run on the diffusion maps to calculate pseudotime, setting the cell with the minimum value in diffusion component 1 as the origin.

*Identifying relationships between ADT and cis-regulatory elements*

Before linking scATAC-seq peaks to putative protein levels, we first ran the SEACells algorithm to increase the robustness of the chromatin data. We ran SEACells using suggested parameters of 75 cells per metacell (120 metacells for the Phospho-seq dataset), n_waypoint_eigs = 10 and waypoint_proportion = 0.9. Based on the SEACells grouping, we calculated cumulative counts values from the ATAC, ADT, and RNA modalities. All three modalities were normalized after aggregation: ADT using CLR normalization, ATAC using TF-IDF and RNA using log-normalization.

To determine candidate peaks associated with ADTs that show regulatory potential for proximal genes, we first found peaks that were correlated to each protein. For this we ran Pearson correlation between each peak and ADT within the metacell object. We first filtered the peaks with a correlation of >0.5, which for SOX2 and OTX2 represents the top ~2.5% of correlated peaks. For ADTs without any peaks with >0.5 correlation, we reduced the correlation cut-off by 0.1 until we observed >1% of peaks correlating. This resulted in a correlation cut-off of 0.4 for TBR2 and 0.3 for pRPS6 and pSTAT3. We next filtered these peaks for those that harbored the corresponding TF binding motif as indicated by the JASPAR database, using a probability score cutoff of >400. Peaks that passed both the correlation-based and sequence-based criteria were linked to the associated ADT.

We next aimed to link these peaks to putative target genes. To do this, we used the LinkPeaks function in Signac with parameters pvalue_cutoff = 0.10, score_cutoff = 0.01. We only considered gene-peak links that fell within 500kb. For each identified gene-peak link, we calculated the Pearson correlation across metacells between peak accessibility and gene expression, and retained all peaks with correlation >0.2 (activating) or <-0.2 (repressive). These cut-offs represented approximately the top 10% positively and negatively correlated genes with ADT expression for each ADT. The lists of each of these peak-gene relationships are in Table S2.

*TF Footprinting*

For TF footprinting analysis, motifs were first identified using the Signac function AddMotifs with JASPAR2020 motif dataset. We only considered motifs within called peaks and then split the peaks into categories based on correlation between peak accessibility and TF protein expression. We selected highly correlated peaks in the top third of expression and a second set of peaks with negative correlation. The Footprint function in Signac was used to sum Tn5 insertion events surrounding motif instances in the highly correlated and negatively correlated peaks. We partitioned cells into ten quantiles based on TF protein expression and displayed Tn5 insertion enrichment for the first and tenth expression quantile using the PlotFootprint function in Signac.

*Chip-Seq Benchmarking*

For benchmarking our SOX2 candidate peaks, we relied upon a previously published SOX2 ChIP-Seq dataset performed on *in vitro* differentiated human neural progenitor cells (hNPCs) [14]. We plotted the ChIP-Seq data alongside our ATAC-seq data by converting the raw data into a BigWig format and plotting with the CoveragePlot function in Signac.

*Gene Ontology Analysis*

The genes associated with the top 500 peak-gene links for each TF were used to analyze gene ontology category enrichment using the enrichr R package [16]. For instances where there were fewer than 500 peak-gene links, all the peaks were used. The GO categories were then filtered to include only those with ≥ five genes as hits.

*STAT3 Active Module Score*

To calculate and plot the STAT3 active module score per cell, we extracted the gene names for all peak-gene links associated with high pSTAT3 signal and increased proximal gene expression. We then used the AddModuleScore() function in Seurat to calculate the collective expression of those genes from the RNA modality in each cell.

### References


1. van Buggenum, J. A. G. L. *et al.* A covalent and cleavable antibody-DNA conjugation strategy for sensitive protein detection via immuno-PCR. *Sci. Rep.* **6**, 22675 (2016).
2. Stoeckius, M. *et al.* Cell Hashing with barcoded antibodies enables multiplexing and doublet detection for single cell genomics. *Genome Biol.* **19**, 224 (2018).
3. Srivatsan, S. R. *et al.* Massively multiplex chemical transcriptomics at single-cell resolution. *Science* **367**, 45–51 (2020).
4. Srivastava, A., Malik, L., Smith, T., Sudbery, I. & Patro, R. Alevin efficiently estimates accurate gene abundances from dscRNA-seq data. *Genome Biol.* **20**, 65 (2019).
5. Stuart, T., Srivastava, A., Madad, S., Lareau, C. A. & Satija, R. Single-cell chromatin state analysis with Signac. *Nat. Methods* **18**, 1333–1341 (2021).
6. Zhang, Y. *et al.* Model-based analysis of ChIP-Seq (MACS). *Genome Biol.* **9**, R137 (2008).
7. Schep, A. N., Wu, B., Buenrostro, J. D. & Greenleaf, W. J. chromVAR: inferring transcription-factor-associated accessibility from single-cell epigenomic data. *Nat. Methods* **14**, 975–978 (2017).

8. Huang, Y., McCarthy, D. J. & Stegle, O. Vireo: Bayesian demultiplexing of pooled single-cell RNA-seq data without genotype reference. *Genome Biol.* **20**, 273 (2019).
9. Giandomenico, S. L., Sutcliffe, M. & Lancaster, M. A. Generation and long-term culture of advanced cerebral organoids for studying later stages of neural development. *Nat. Protoc.* **16**, 579–602 (2020).
10. Choudhary, S. & Satija, R. Comparison and evaluation of statistical error models for scRNA-seq. *Genome Biol.* **23**, 27 (2022).
11. Hao, Y. *et al.* Dictionary learning for integrative, multimodal, and scalable single-cell analysis. Preprint at https://doi.org/10.1101/2022.02.24.481684.
12. Angerer, P. *et al. destiny*: diffusion maps for large-scale single-cell data in R. *Bioinformatics* vol. 32 1241–1243 Preprint at https://doi.org/10.1093/bioinformatics/btv715 (2016).
13. Cao, J. *et al.* The single-cell transcriptional landscape of mammalian organogenesis. *Nature* **566**, 496–502 (2019).
14. Zhou, C. *et al.* Comprehensive profiling reveals mechanisms of SOX2-mediated cell fate specification in human ESCs and NPCs. *Cell Res.* **26**, 171–189 (2016).
15. Quinlan, A. R. & Hall, I. M. BEDTools: a flexible suite of utilities for comparing genomic features. *Bioinformatics* **26**, 841–842 (2010).
16. Kuleshov, M. V. *et al.* Enrichr: a comprehensive gene set enrichment analysis web server 2016 update. *Nucleic Acids Res.* **44**, W90–7 (2016).
